# Computational Evaluation of Phytochemicals as Potential Anti-HIV Drugs Targeting CCR5 and CXCR4 Receptors

**DOI:** 10.1101/2025.04.10.648243

**Authors:** Sadman Sakib Nebir, Tawsif Al Arian, Bishajit Sarkar, Ripa Moni, Salina Malek, Syed Sajidul Islam, Umme Salma Zohora, Mohammad Shahedur Rahman

**Affiliations:** Department of Pharmacology, Faculty of Basic Sciences, Bangladesh University of Health Sciences, Dhaka-1216, Bangladesh; Department of Pharmacy, Jahangirnagar University, Dhaka-1342, Bangladesh; Department of Biotechnology and Genetic Engineering, Faculty of Biological Sciences, Jahangirnagar University, Dhaka-1342, Bangladesh

**Keywords:** Antiviral agents, CCR5, CXCR4, Drug design, HIV, Phytochemicals

## Abstract

HIV is a major worldwide health concern; hence new therapeutic approaches are needed to fight viral resistance and enhance treatment results. HIV entrance into host cells depends on the CCR5 and CXCR4 receptors, which makes them potential targets for antiviral medication development. The objective of this study is to computationally evaluate 53 phytochemicals that target CCR5 and CXCR4 as potential anti-HIV medications. Effective anti-HIV medications were projected to be phytochemicals that may inhibit these receptors and so interfere with the HIV life cycle. AutoDock Vina was used to perform the molecular docking investigation from which six phytochemicals capable of inhibiting CCR5 and CXCR4 were identified based on the lowest docking score. i.e., Withaferin A, Oleanolic Acid, Ursolic Acid, Theaflavine, Camptothecin, and Hypericin. The SWISSADME server was utilized to decide their druglikeness properties, the ADMETlab server to predict different pharmacokinetic and pharmacodynamic properties, the PASS-Way2Drug server to evaluate their activity spectra, and the RS-WebPredictor server to figure out the metabolism in the body. They adhered to Lipinski’s rule of five and had promising ADME/toxicity study result along with favorable molecular dynamics simulation. Overall, the above-mentioned six phytochemicals might have the potential to be used as alternative HIV therapeutics.

## 1. Introduction

Human immunodeficiency virus (HIV) raids the body’s immune system and weakens a person’s defenses against many infections and some cancers that people with healthy immune systems can fight more easily. Infected individuals eventually lose their immunological protection and become immune-deficient as the virus kills and damages immune cells. CD4 cell count is commonly used to assess immune function (1). Approximately 40.1 million [33.6-48.6 million] people have died from HIV since the epidemic’s inception, while 84.2 million [64.0-113.0 million] individuals have contracted the infection with HIV. At the end of 2021, there were 38.4 million [33.9-43.8 million] HIV-positive individuals worldwide. Moreover, 650000 people died of HIV-related illnesses worldwide in 2021 (2). Within the family Retroviridae, subfamily Orthoretrovirinae, the genus Lentivirus contains the human immunodeficiency virus (HIV) (3). Two identical single-stranded RNA molecules enclosed inside the virus particle’s core comprise the HIV genome (3). HIV is a retrovirus. The genetic material is single-stranded RNA, not the double-stranded DNA found in human cells. Retroviruses also have reverse transcriptase, which can convert RNA into DNA and infect humans or host cells with that DNA. When HIV infects a cell, it first binds to the host cell and unites with it. The virus then uses the host cell mechanism to convert viral RNA into DNA and multiply. The new HIV copies then leave the host cell and spread to other cells. Binding and invasion into host cells are the first step in the human immune deficiency virus (HIV) replication cycle, which determines the tropism of the virus and the potential for HIV to impair the human immune system. HIV evades the host immune response by delivering its DNA into the host cell’s cytoplasm through a multi-step process (4). In the first phase of the viral replication cycle, HIV entry starts with the virus adhering to the host cell and ends with the cell and viral membranes fusing and the viral core being delivered into the cytoplasm (4). The second step in viral entry and essential for infection is that the envelope (Env) protein binds to the host cell protein CD4, the primary receptor. HIV envelope (Env) protein is comprised of gp41 and gp120 subunits (5, 6), and Env is a highly glycosylated trimmer of the gp41 and gp120 heterodimers. Receptor binding is carried out by the gp120 subunit, which has five relatively conserved domains (C1–C5) and five variable loops (V1–V5), so termed for their relative genetic heterogeneity (4). CD4, an immunoglobulin superfamily member, typically enhances T-cell receptor (TCR) mediated signaling (7). We will focus on the third step of virus entry: co-receptor CCR5 (C-C chemokine receptor type 5) and CXCR4 (C-X-C chemokine receptor type four) binding. Various HIV strain uses different co-receptors. R5 HIV uses the CCR5 chemokine receptor, X4 HIV uses CXCR4, and R5X4 HIV uses both co-receptors (8). There is no convincing evidence that co-receptors other than CCR5 and CXCR4 are important in supporting HIV-1 infection in vivo. With rare exceptions, only the R5 and R5X4 viruses are typically spread from person to person (9). Chemokine receptors are crucial during the initial stages of CD4+ cells’ HIV-1 infection. While CCR5 permits entry of M-tropic HIV-1 strains, CXCR4 is the fusogenic receptor that favors entry of T-tropic HIV-1 strains (10). The fourth step in the virus entry process is moving the viral particle to where productive membrane fusion occurs. Recent research has demonstrated that several viruses usurp cellular transport systems to reach specific locations that are either required for infection or efficient entrance, and HIV may also do the same to reach locations where membrane fusion can take place (11–13). The chemokine receptor CCR5 is expressed on various cell types, including epithelial, endothelial, vascular smooth muscle, and fibroblasts; dendritic cells, macrophages, and memory T cells of the immune system; neurons, microglia, and astrocytes of the central nervous system (14). It was discovered in 1996 that CCR5 is required as a co-receptor for entry of the macrophage tropic HIV strains (15, 16). CCR5 ligands impede viral entry and envelope-mediated fusion (15). The virus prefers CCR5 during initial infection, whereas the alternative co-receptor ‘C-X-C chemokine receptor type four’ (CXCR4) is utilized considerably later in HIV infection, as the infected person approaches AIDS (17). Neutralizing antibodies, genetic modification, and preventing an integrated latent provirus are some of the fundamentally novel methods being developed for treating HIV-1. Nowadays, most medications target one of the HIV-1 enzymes, such as reverse transcriptase, protease, or integrase. NRTIs (Nucleoside and nucleotide reverse transcriptase inhibitors) and NNRTIs (non-nucleoside reverse transcriptase inhibitors) are the two categories into which reverse transcriptase inhibitors are typically divided (18). Protease inhibitors are a second significant class of clinically utilized inhibitors. In addition to inhibitors of the HIV-1 enzyme, inhibitors that affect other steps in the viral life cycle are being developed. Viral cell entry inhibitors used in HIV infection can be divided into inhibitors of viral-cell membrane fusion and inhibitors of viral envelope proteins’ ability to attach to receptors (18). The G-protein coupled chemokine receptors CCR5 and CXCR4 determine the cellular tropism of HIV-1. The primary cell infection is specifically caused by viruses that preferentially employ CCR5 (R5 strains), as this co-receptor is mostly found in macrophages. On the other hand, viruses’ ability to replicate themselves is due to their preference for interacting with CXCR4 (X4 strains). This second co-receptor is mostly expressed in T-lymphocytes (19). These findings imply that CCR5 is a good target for HIV prevention or treatment and that even novel strategies, like anti-CCR5 vaccination, can offer vital scientific understanding and, more importantly, effective tools for the battle against HIV and other immune-based diseases. It has been demonstrated that several types of phytochemicals prevent HIV replication, and various phytochemicals that target the HIV infection cycle are investigated here. Natural products originating from plants continue to be a source for developing novel medicines, such as anti-HIV agents. Despite significant advances in HIV chemotherapy, there is still a need for novel anti-HIV drug development, and medicinal plants can be beneficial in this effort. Several plant species, particularly *Artemisia annua*, *Garcinia edulis*, *Justicia gendarussa*, *Phyllanthus pulcher*, *Rhus chinensis*, *Smilax corbularia*, *Terminalia paniculata*, and *Tuberaria lignosa*, have demonstrated notable anti-HIV activity (20). As a result, the use of plant-based alternatives holds enormous promise for the treatment of HIV patients. Now, the focus is shifting to using natural compounds, such as plant-based alternatives, to produce anti-HIV medications. Although HIV/AIDS is not a new disease, clinical trials are currently being conducted on research based on plant-derived products. Therefore, this computational study aimed to analyze antiviral phytochemicals as potential treatment options for HIV.

**Table 1.** List of the plant-derived anti-HIV agents that work via CCR5 and CXCR4.

| Serial | Name of the agents | PubChem CID | Reference |
| --- | --- | --- | --- |
| 1 | Ellagic acid | 5281855 | (21) |
| 2 | Gallic acid | 370 | (21) |
| 3 | Epigallocatechin gallate | 65064 | (22) |
| 4 | Quercetin | 5280343 | (23) |
| 5 | Quercetrin | 5280459 | (23) |
| 6 | Indirubin | 10177 | (24) |
| 7 | Fisetin | 5281614 | (25) |
| 8 | Naringenin | 439246 | (26) |
| 9 | Luteolin | 5280445 | (27) |
| 10 | Apigenin | 5280443 | (28) |
| 11 | Camptothecin | 24360 | (29) |
| 12 | Daidzein | 5281708 | (30) |
| 13 | Kaempferol | 5280863 | (31) |
| 14 | 6-Gingerol | 442793 | (32) |
| 15 | Curcumin | 969516 | (33) |
| 16 | Palmatine | 19009 | (34) |
| 17 | Berberine | 2353 | (35) |
| 18 | Magnoflorine | 73337 | (36) |
| 19 | Lanuginosine | 97622 | (37) |
| 20 | Liriodenine | 10144 | (38) |
| 21 | Anonaine | 160597 | (39) |
| 22 | Ursolic acid | 64945 | (40) |
| 23 | Eugenol | 3314 | (41) |
| 24 | Cordycepin | 6303 | (42) |
| 25 | Harmaline | 3564 | (43) |
| 26 | Harmine | 5280953 | (44) |
| 27 | Lycorine | 72378 | (45) |
| 28 | Oliverine | 24198103 | (46) |
| 29 | Hesperetin | 72281 | (47) |
| 30 | Oleanolic acid | 10494 | (48) |
| 31 | Myricetin | 5281672 | (49) |
| 32 | Imperatorin | 10212 | (50) |
| 33 | Bergapten | 2355 | (51) |
| 34 | Silybin | 31553 | (52) |
| 35 | Withaferin A | 265237 | (53) |
| 36 | Cordioside | 101915817 | (54) |
| 37 | Neoandrographolide | 9848024 | (55) |
| 38 | Rotenone | 6758 | (56) |
| 39 | Calanolide A | 64972 | (57) |
| 40 | Pericalline | 6436240 | (58) |
| 41 | Ferruginol | 442027 | (59) |
| 42 | Allicin | 65036 | (60) |
| 43 | Genistein | 5280961 | (61) |
| 44 | Scutellarein | 5281697 | (62) |
| 45 | Capsaicin | 1548943 | (63) |
| 46 | Theaflavin | 135403798 | (64) |
| 47 | Hypericin | 3663 | (65) |
| 48 | Methyl rosmarinate | 6479915 | (66) |
| 49 | Licoleafol | 11111496 | (67) |
| 50 | Nimocinol | 178770 | (68) |
| 51 | Isomeldenin | 76316558 | (69) |
| 52 | Tembetarine | 167718 | (70) |
| 53 | Tinosponone | 15215479 | (71) |

**Figure 1.**
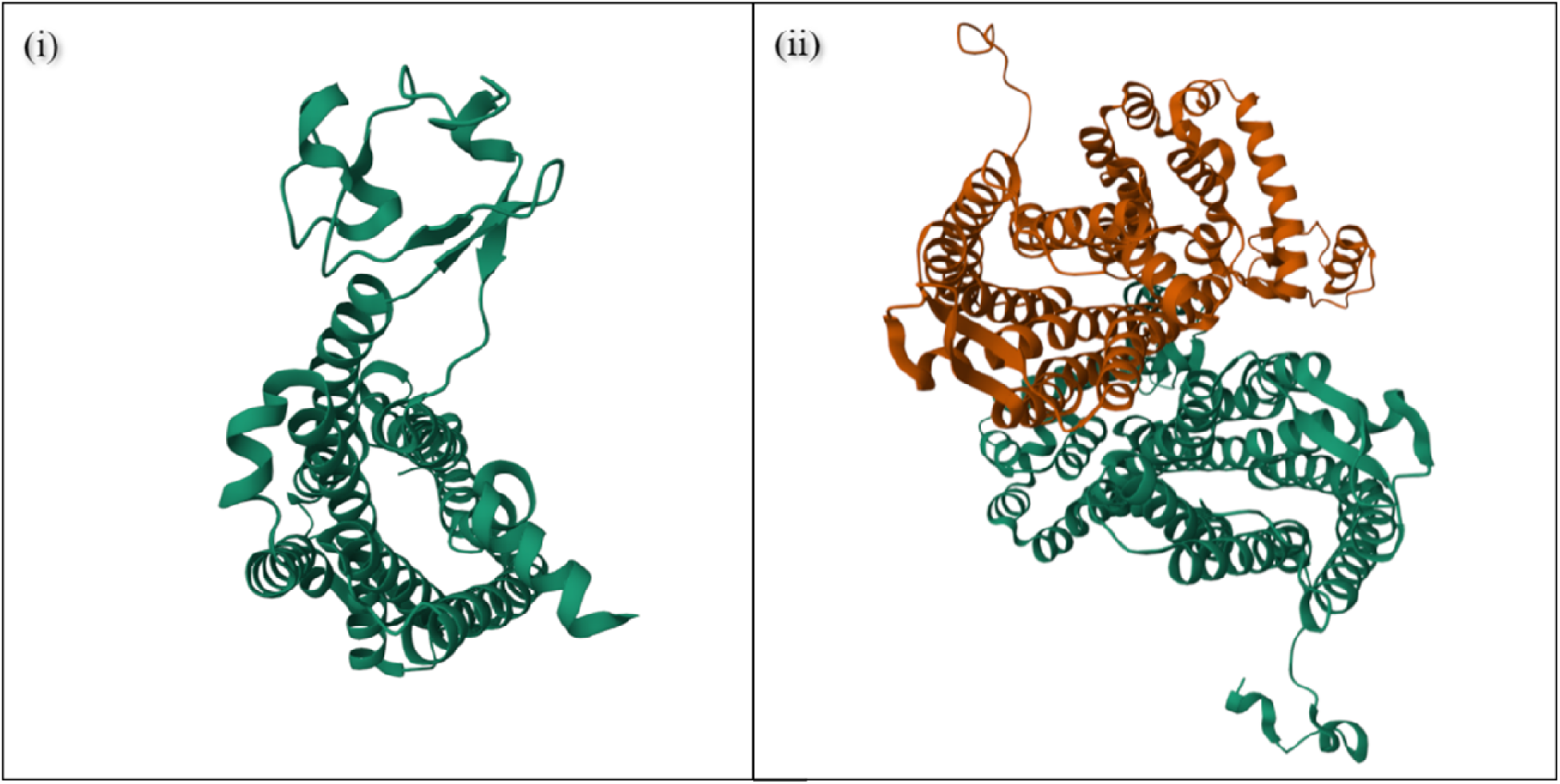
The 3D structures of the target proteins: 1. CCR5 Chemokine Receptor (PDB ID: 4MBS) (72); 2. CXCR4 chemokine receptor in complex (PDB ID: 3ODU) (73). The 3D structures were visualized by Protein Data Bank 3D viewer.

### 1.1 Role of CCR5 and CXCR4 in terms of HIV infection

Immune cells, such as CD4+ T-cells, essential in the immunological response against viruses, bacteria, and other pathogens, have the protein CCR5 on their surface. CCR5, CCR5, also known as CCCKR-5 or CKR5, is engaged mainly in immune surveillance, inflammatory response, tumor development and metastasis (20)(72)(73), etiology of inflammatory disorders (74)(75)(76), asthma (77)(78), and cancer (72, 73). It is vital in the recruitment of immune cells to inflammatory sites by guiding immune cell movement (chemotaxis) along the chemokine gradient (79) (80). CCR5 modulates the trafficking and effector activities of memory/effector T lymphocytes, macrophages, and immature dendritic cells (81). Aside from its direct role in immune process mediation, it also suppresses learning, memory, and synaptic connections in the brain (82). The human immunodeficiency virus (HIV) uses CCR5 as a co-receptor in order to enter and infect CD4+ T-cells. CCR5 is a seven-transmembrane G protein-coupled receptor (GPCR) that belongs to the class A GCPR family. CCR5 is a GPCR with seven transmembran a-helices, three extracellular loops, three intracellular loops, an amino-terminal domain, and a carboxyl-terminal domain (83).

HIV is an RNA virus that attaches to CD4 on the surface of target cells through the surface glycoprotein gp120. HIV cannot enter the cell through this contact; it needs a co-receptor, CCR5 or CXCR4, to finish the entrance process. The majority of HIV strains, sometimes referred to as R5-tropic HIV, employ CCR5 as their co-receptor. Natural CCR5 ligands contain chemokines (small chemoattractant cytokines) involved in innate immunity, which are natural suppressors of HIV-1 infection (84–87): macrophage inflammatory proteins CCL3 (MIP-1 a) and CCL4 (MIP-1 b), CCL5 (RANTES - regulated on activation, normal T-cell expressed and secreted), and CCL3L1, the most potent among CCR5 (88). CCL7 (MCP-3) is the main antagonist ligand of the CCR5 receptor (89). The CCR5 receptor is activated by its agonist ligands, which increase cell migration and mediate inflammatory reactions. In summary, CCR5 is a co-receptor essential for HIV entrance into target cells (Figure 3).

**Figure 2.**
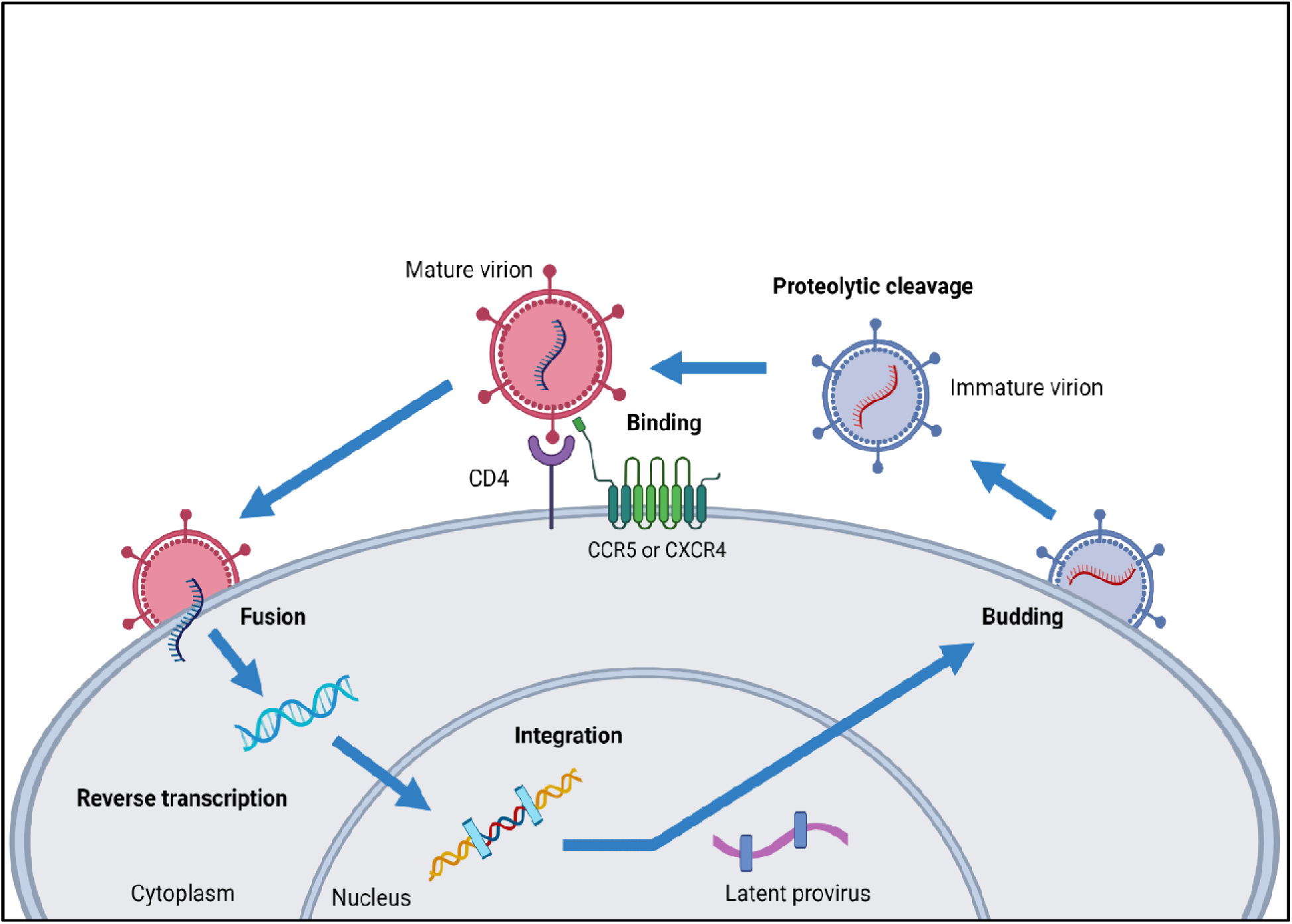
Extracellular virions penetrate their target cell through a three-step process: binding to the CD4 receptor, binding to the CCR5 or CXCR4 coreceptors, or both, and membrane fusion. The HIV reverse transcriptase enzyme catalyzes HIV RNA transcription into double-stranded HIV DNA, a phase blocked by nucleoside analogs and non-nucleoside reverse transcriptase inhibitors (NNRTIs). HIV integrase is an enzyme that aids in integrating HIV DNA into host chromosomes. Immature virions are created and bud from the cell surface following transcription and translation of the HIV genome. Since the HIV protease enzyme cleaves polypeptide chains, the virus can mature.

**Figure 3:**
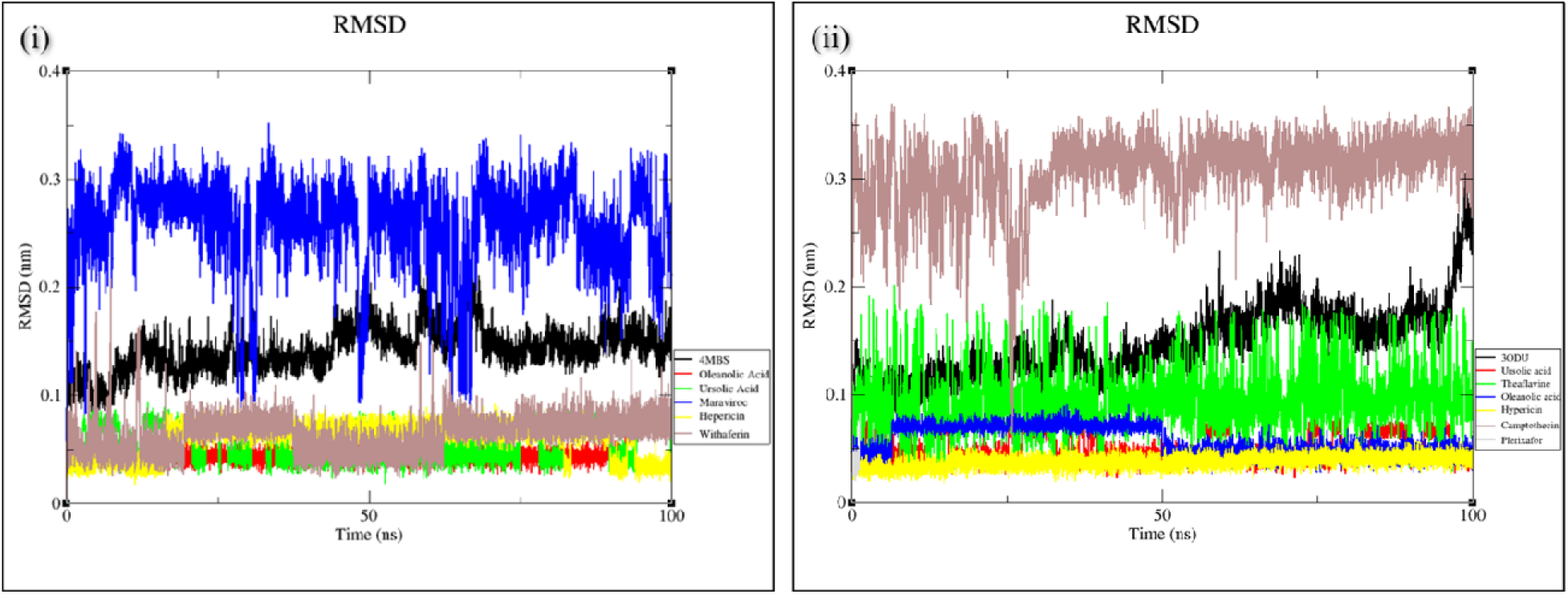
RMSD (Root Mean Square Deviation) of (i) Protein RCSB ID: 3ODU (Black color) and Ligand(s): Ursolic Acid (Red), Theaflavin (Green), Oleanolic Acid (Blue), Hypericin (Yellow), Camptothecin (Brown), Plerixafor (Gray) (ii) Protein RCSB ID: 4MBS (Black) and Ligand(s): Oleanolic Acid (Red), Ursolic acid (Green), maraviroc (Blue), Hypericin (Yellow), Withaferin A (Brown)

CXCR4, commonly known as “fusin,” is one of the most well-studied chemokine receptors due to its previously discovered role as an HIV entrance coreceptor (87). Although CXCR4 is known to bind only CXCL12, another chemokine receptor, CXC receptor 7 (CXCR7, ACKR3, RDC1, CMKOR1, or GPR159), was identified in 2005 as a CXCL12 receptor (90). Further research will be necessary to determine the precise involvement of the CXCR4-CXCR7-CXCL12 axis in cell migration (87). Certain chemokines, including MIP-1a, MIP-1b, and RANTES, were identified to suppress HIV-1 infection, indicating that the chemokine receptors may be a coreceptor for HIV-1 entrance (91). It has been demonstrated to have an essential role in the pathogenesis of HIV infection. Soon after CCR5, the a-chemokine receptor CXCR4 was identified as the predominant coreceptor for T-tropic HIV-1 isolates that can replicate successfully in T-cell lines and primary CD4+ T cells (88).

CXCR4 and CCR5 chemokine receptors are coreceptors for HIV-1 entrance into CD4+ cells. At the beginning stages of HIV infection, viral segregates frequently utilize CCR5, while later disconnects typically use CXCR4. The pattern of expression and regulation of these chemokine receptors on T cell subsets has crucial implications for AIDS pathogenesis and lymphocyte recirculation.

## 2. Materials and Methods

### 2.1 Target Selection and Preparation

In this work, the CCR5 Chemokine Receptor (4MBS) and the CXCR4 Chemokine Receptor (3ODU) were chosen as possible targets. PDB structures of targets with bound ligands were obtained (Protein Data Bank). Structures lacking the physiologically required cofactor and heteroatom were rejected, as were those lacking side chains. By preserving the optimum physiological pH, hydrogen atoms were introduced to the target. Apart for those essential to the aim hypothesis, all water molecules were eliminated. The Kollman Charges were used to determine the partial charges. Nonpolar hydrogens were merged since it is a mechanism for representing unified atoms. AutoGrid was used to generate grid maps, which display a 3D grid of interaction energies with the target.

### 2.2 Ligand Selection and Preparation

Potential ligands were identified and obtained from PubChem using 3D SDF format structures. As AutoDock Vina cannot read the SDF format, the PyMOL tool was used to convert it to PDB format. All of the PDB files were then converted to PDBQT files and loaded into AutoDock in order to dock with the required targets. Positive control for 4MBS and 3ODU, Maraviroc (Positive control, PubChem CID 3002977), and Plerixafor (Positive control, PubChem CID 65015), were prepared using the same process.

### 2.3 Receptor Grid Generation

Grid usually confines the active site to shorten specific areas of the receptor protein so that the ligand can dock specifically. In Glide, a grid was generated using the default Van der Waals radius scaling factor of 1.0 and charge cutoff of 0.25. The spacing and npts were set to a default of 0.375 and 40. The energy range and exhaustiveness were 4 and 8, the default value. A cubic box is available (reference ligand active site). Then, the grid box volume was adjusted to 168.307× 116.536× 39.126 for the docking test.

### 2.4 Pass Prediction

PASS-Way2Drug (http://way2drug.com/passonline/) online server was used to predict the PASS (Prediction of Activity Spectra for Substances) values of the best selected phytochemicals or ligands (74). This server can predict over 4000 different types of biological, pharmacological, metabolic, pharmacokinetic, and toxic characteristics of chemical compounds by analyzing its database consisting of 30000 biologically active compounds (75). The canonical smiles of these phytochemicals were used in the PASS analysis, and the probability of a compound being active or the Pa threshold was kept greater than 0.7 or 70% to increase the reliability of the predictions. Thereafter, SMARTCyp (https://smartcyp.sund.ku.dk/mol_to_som) online server was used to carry out the P450 site of metabolism or SOM prediction (76). The server utilizes the following standard equation for conducting the predictions:

Score, S = E - 8*A - 0.04*SASA

Where S represents the prediction score, E represents the activation energy score, A is the accessibility measure of the compound in the query determined based on its topological features, and SASA is the Solvent Accessible Surface Area that is also measured from the topological characteristics of the query compound. The lower the prediction score and energy score are, the better the reliability of the prediction is, and vice versa (45). Canonical smiles of the phytochemical were again used in the server to perform the predictions.

### 2.5 Ligand-based drug-likeness property analysis and ADME/toxicity prediction

SWISSADME server (http://www.swissadme.ch/) was used to assess the drug-likeness properties of the nine selected ligand molecules (95). In order to predict their various pharmacokinetic and pharmacodynamic properties, the ADMET analysis for each of the ligand molecules was carried out using online-based servers, admetSAR (http://lmmd.ecust.edu.cn/admetsar2/) and ADMETlab (http://admet.scbdd.com/) (96, 97). Both the servers determined the absorption, distribution, metabolism, excretion, and toxicity properties. The numeric and categorical values of the results given by the ADMETlab server were changed into qualitative values according to the explanation and interpretation described in the ADMETlab server (http://admet.scbdd.com/home/interpretation/) for the convenience of interpretation.

### 2.6 Molecular Dynamic Simulation

Molecular dynamics is a sophisticated automated simulation approach used for evaluating the degree of stability of protein and protein-ligand complex structure at the minuscule stage through demonstrating the behavioral characteristics, interacting arrangement, fluctuation, physical foundation of function, and structure. The molecular dynamics (MD) simulations were conducted using GROMACS version 2023.1 to investigate the behavior of both Apo protein and Protein-ligand complexes (77). The protein was parameterized for its protein content using the CHARMM General Force Field. The SwissParam server was utilized to perform the ligand topologies (78). The structures underwent 2500 cycles of vacuum minimization using the steepest descent method in order to mitigate any potential steric issues. The solvation of the structure was accomplished through the utilization of the Simple Point Charge (SPC) water model. Subsequently, the system was rendered neutral through the introduction of Na+ and Cl-ions utilizing the gmx genion tool. This measure was implemented in order to maintain the overall electrical neutrality of the system. After minimization, three stages were led in the MD simulation: NVT, NPT, and production. The equilibration of the systems was conducted in two phases. Initially, an NVT equilibration lasting 100 picoseconds was conducted to attain a steady state of the number of particles, volume, and temperature. The purpose of this step was to elevate the system to a temperature of 300 Kelvin. The second step involved conducting a 100 picosecond NPT equilibration, which aimed to maintain the system’s pressure and density stability by ensuring an equal number of particles, pressure, and temperature. The simulations involved the imposition of bond constraints on all bonds within the protein, thereby inducing position restraint of the protein group. The constrained conditions of NVT and NPT resulted in the relaxation of water molecules surrounding the protein, leading to a decrease in system entropy. The dynamics were conducted utilizing the Parrinello-Rahman barostat and the v-rescale thermostat (79). The relaxation of the barostat and thermostat persisted for a duration of 100 picoseconds. The application of Linear Constraint Solver algorithm was used to impose constraints on the covalent bonds. The Particle-Mesh Ewald (PME) method was utilized to process the electrostatic interactions. After reaching equilibrium, every framework underwent a production run that lasted for 100 nanoseconds (ns) of simulation time.

## 3. Results and Discussion

### 3.1 Molecular Docking

The study conducted molecular docking of identified ligand molecules with their receptor proteins, finding that molecules with the least binding energy were most effective in hindering their receptors due to their expanded binding affinity (Table 2) (89). Among the ligands, Ursolic acid, Oleanolic acid, Withaferin A, Hypericin (inhibit CCR5) and Camptothecin, Ursolic acid, Oleanolic acid, Theaflavin, and Hypericin (inhibit CXCR4), were chosen for their lower free binding energy and comparable binding energy to respective controls Maraviroc and Plerixafor (table 2). An efficient method for computer-aided drug design is molecular docking, which uses particular algorithms to assign affinity scores based on the positions of ligands inside target binding pockets. Since the complex will spend more time in contact, the lowest docking score indicates the highest affinity (90, 91). Strengthening the receptor-ligand interactions requires both hydrophobic and hydrogen interactions (92). Multiple hydrogen and hydrophobic interactions were predicted to be formed by the six best-selected ligands and the other ligands with the target molecules (Figure 9 and Table 2 & 12). Therefore, Ursolic acid, Oleanolic acid, Withaferin A, and Hypericin, Camptothecin, and Theaflavin ligands from each receptor category were chosen and assessed in the subsequent phases of the study.

**Table 2.**
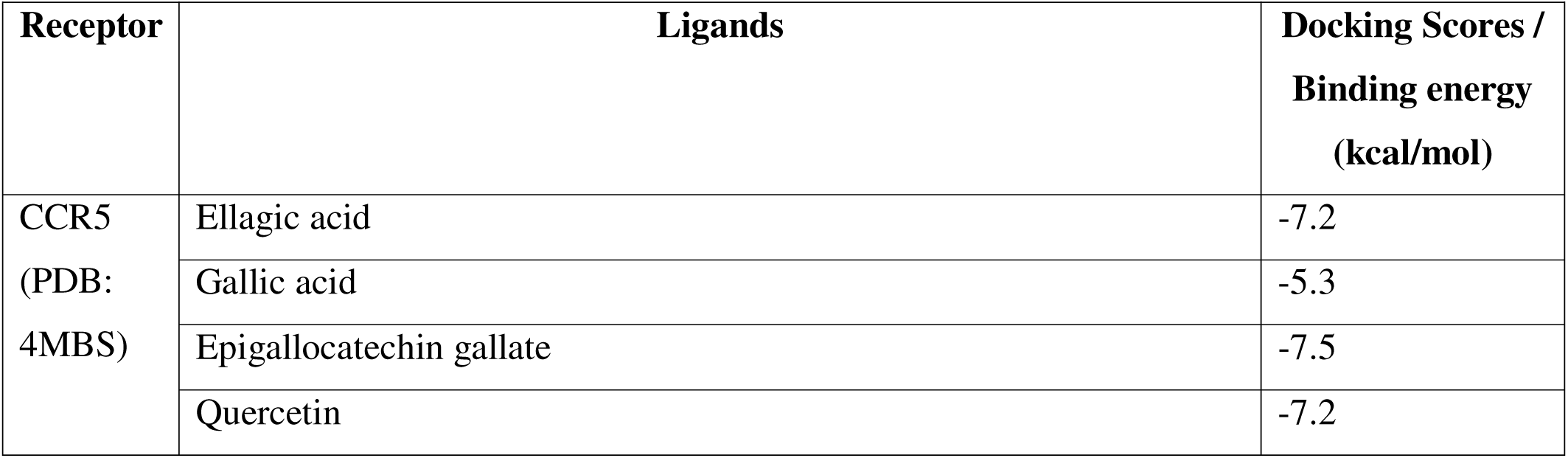
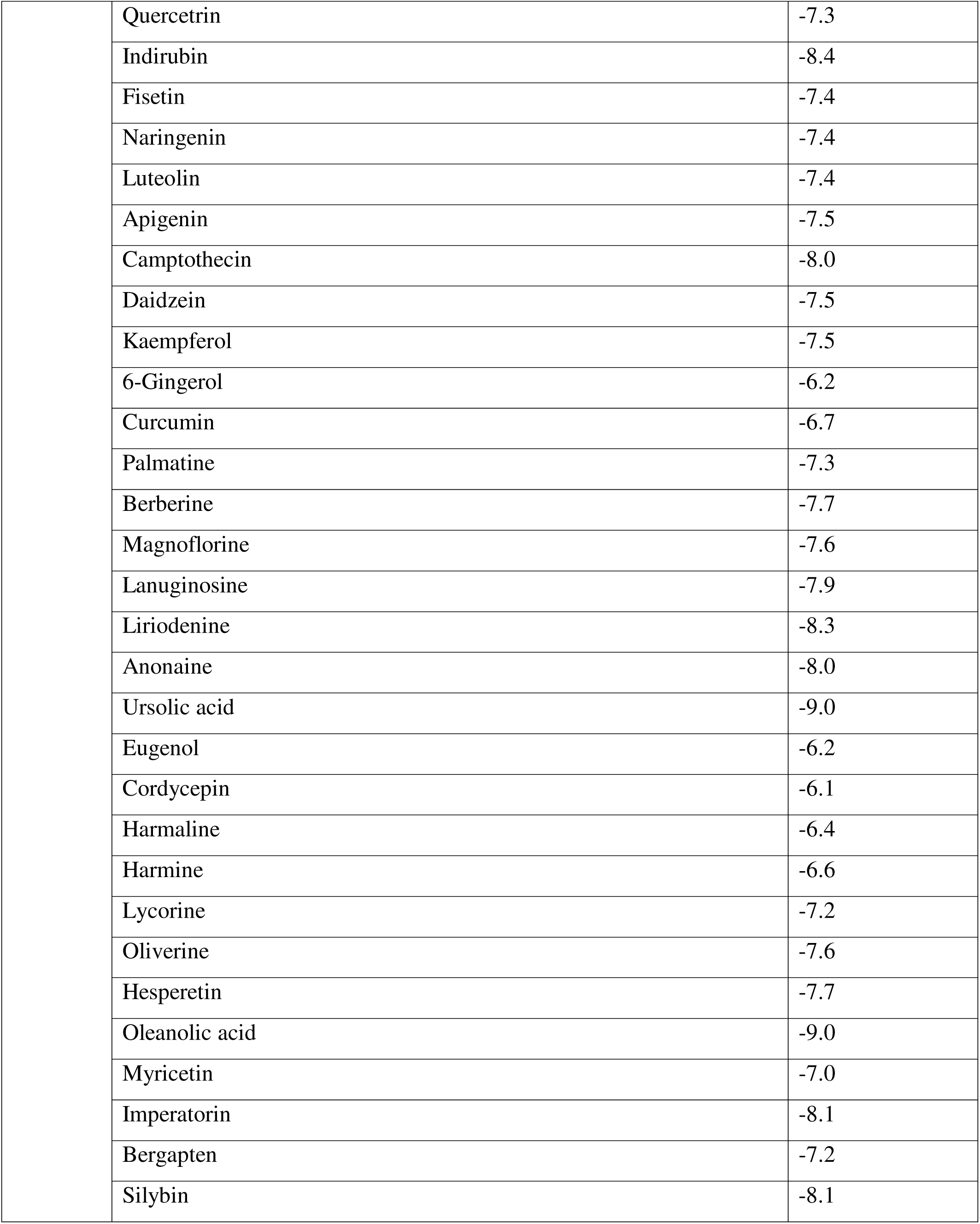
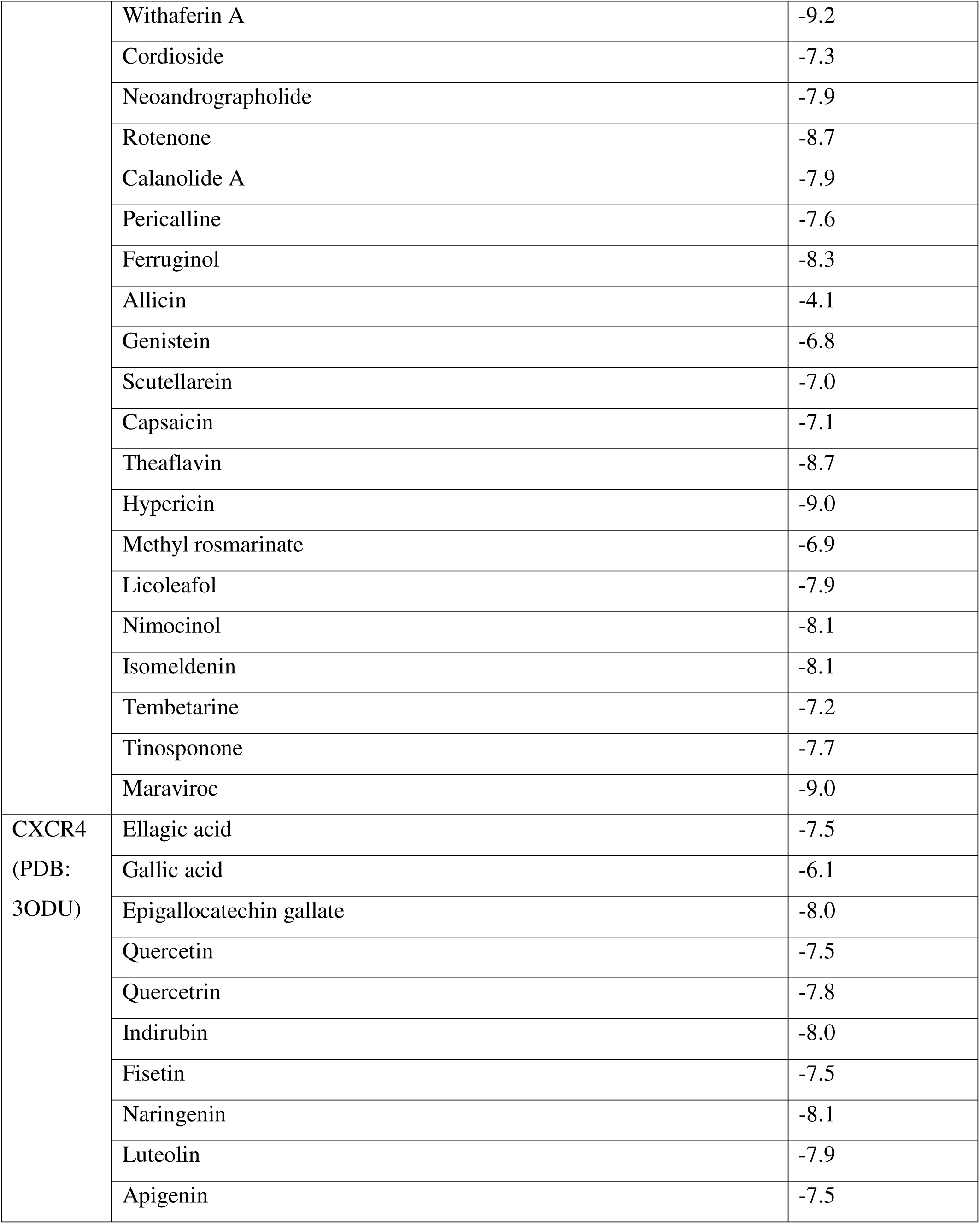
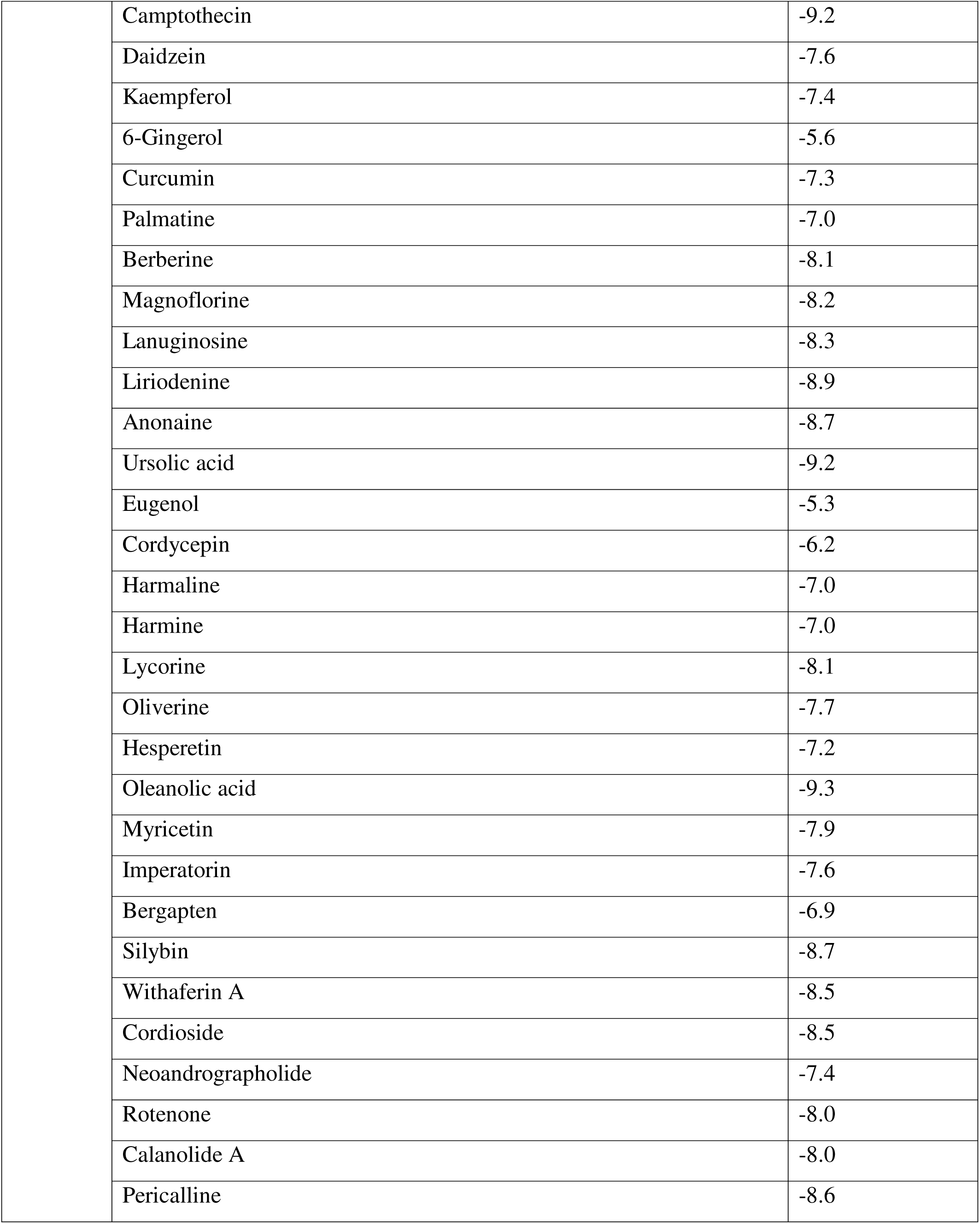
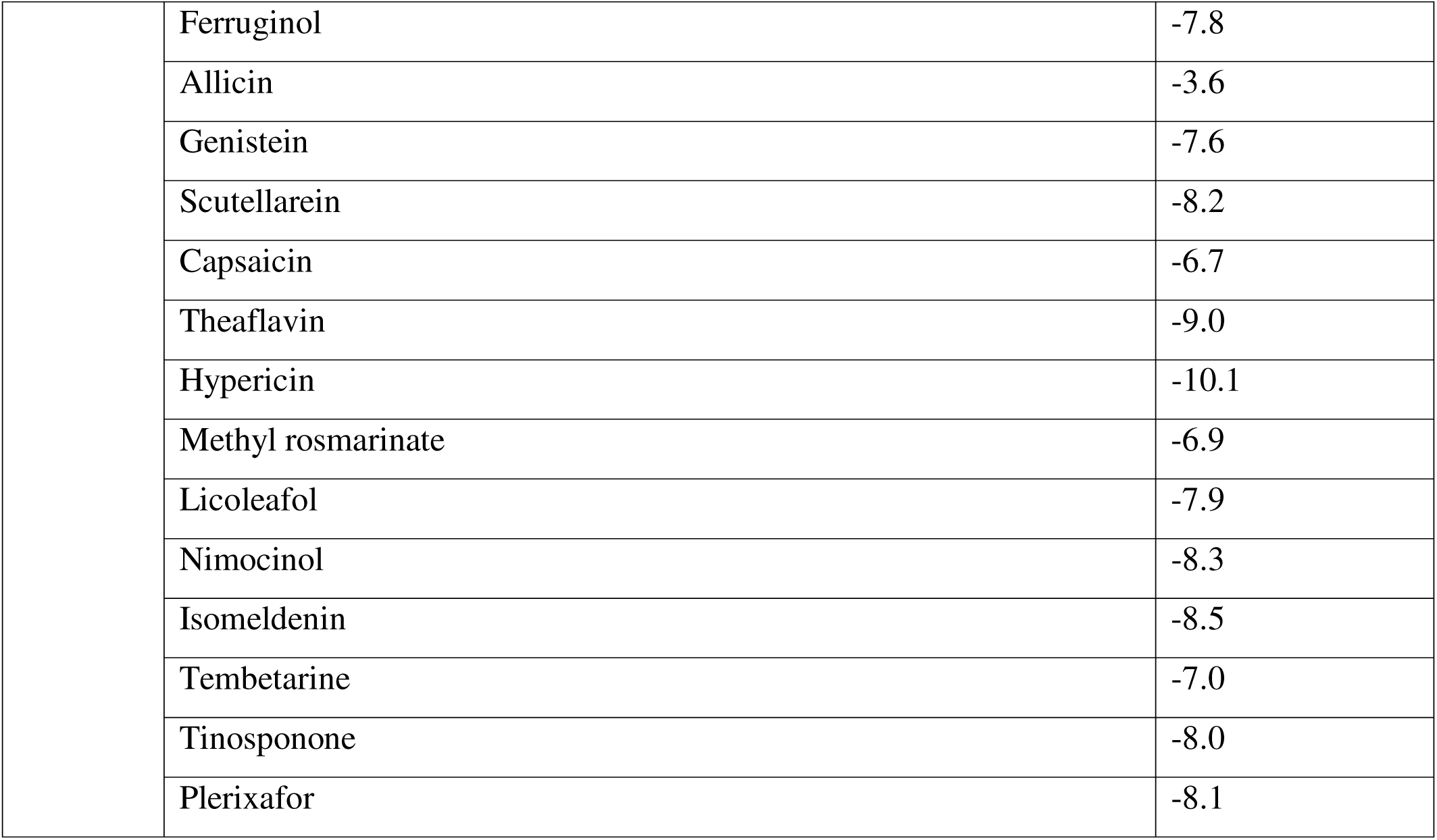
Docking Scores.

| Receptor | Ligands | Docking Scores / Binding energy (kcal/mol) |
| --- | --- | --- |
| CCR5<br>(PDB: 4MBS) | Ellagic acid | -7.2 |
|  | Gallic acid | -5.3 |
|  | Epigallocatechin gallate | -7.5 |
|  | Quercetin | -7.2 |

|  |  |
| --- | --- |
| Quercetrin | -7.3 |
| Indirubin | -8.4 |
| Fisetin | -7.4 |
| Naringenin | -7.4 |
| Luteolin | -7.4 |
| Apigenin | -7.5 |
| Camptothecin | -8.0 |
| Daidzein | -7.5 |
| Kaempferol | -7.5 |
| 6-Gingerol | -6.2 |
| Curcumin | -6.7 |
| Palmatine | -7.3 |
| Berberine | -7.7 |
| Magnoflorine | -7.6 |
| Lanuginosine | -7.9 |
| Liriodenine | -8.3 |
| Anonaine | -8.0 |
| Ursolic acid | -9.0 |
| Eugenol | -6.2 |
| Cordycepin | -6.1 |
| Harmaline | -6.4 |
| Harmin | -6.6 |
| Lycorine | -7.2 |
| Oliverine | -7.6 |
| Hesperetin | -7.7 |
| Oleanolic acid | -9.0 |
| Myricetin | -7.0 |
| Imperatorin | -8.1 |
| Bergapten | -7.2 |
| Silybin | -8.1 |
|  | Withaferin A | -9.2 |
|  | Cordioside | -7.3 |
|  | Neoandrographolide | -7.9 |
|  | Rotenone | -8.7 |
|  | Calanolide A | -7.9 |
|  | Pericalline | -7.6 |
|  | Ferruginol | -8.3 |
|  | Allicin | -4.1 |
|  | Genistein | -6.8 |
|  | Scutellarein | -7.0 |
|  | Capsaicin | -7.1 |
|  | Theaflavin | -8.7 |
|  | Hypericin | -9.0 |
|  | Methyl rosmarinate | -6.9 |
|  | Licoleafol | -7.9 |
|  | Nimocinol | -8.1 |
|  | Isomeldenin | -8.1 |
|  | Tembetarine | -7.2 |
|  | Tinosponone | -7.7 |
|  | Maraviroc | -9.0 |
| CXCR4<br>(PDB:<br>3ODU) | Ellagic acid | -7.5 |
|  | Gallic acid | -6.1 |
|  | Epigallocatechin gallate | -8.0 |
|  | Quercetin | -7.5 |
|  | Quercetrin | -7.8 |
|  | Indirubin | -8.0 |
|  | Fisetin | -7.5 |
|  | Naringenin | -8.1 |
|  | Luteolin | -7.9 |
|  | Apigenin | -7.5 |

|  |  |
| --- | --- |
| Camptothecin | -9.2 |
| Daidzein | -7.6 |
| Kaempferol | -7.4 |
| 6-Gingerol | -5.6 |
| Curcumin | -7.3 |
| Palmatine | -7.0 |
| Berberine | -8.1 |
| Magnoflorine | -8.2 |
| Lanuginosine | -8.3 |
| Liriodenine | -8.9 |
| Anonaine | -8.7 |
| Ursolic acid | -9.2 |
| Eugenol | -5.3 |
| Cordycepin | -6.2 |
| Harmaline | -7.0 |
| Harmin | -7.0 |
| Lycorine | -8.1 |
| Oliverine | -7.7 |
| Hesperetin | -7.2 |
| Oleanolic acid | -9.3 |
| Myricetin | -7.9 |
| Imperatorin | -7.6 |
| Bergapten | -6.9 |
| Silybin | -8.7 |
| Withaferin A | -8.5 |
| Cordioside | -8.5 |
| Neoandrographolide | -7.4 |
| Rotenone | -8.0 |
| Calanolide A | -8.0 |
| Pericalline | -8.6 |
| Ferruginol | -7.8 |
| Allicin | -3.6 |
| Genistein | -7.6 |
| Scutellarein | -8.2 |
| Capsaicin | -6.7 |
| Theaflavin | -9.0 |
| Hypericin | -10.1 |
| Methyl rosmarinate | -6.9 |
| Licoleafol | -7.9 |
| Nimocinol | -8.3 |
| Isomeldenin | -8.5 |
| Tembetarine | -7.0 |
| Tinosponone | -8.0 |
| Plerixafor | -8.1 |

### 3.2 Drug-likeness properties

Druglikeness properties of a chemical compound determine the ability of that compound to act as a drug in humans based on its biological and pharmacological properties. Lipinski’s rule of five is a set of widely used rules that are used to evaluate a chemical’s drug-likeness properties. According to Lipinski’s rule of five, an active drug molecule should violate no more than one of these primary criteria: Molecular weight <500 g/mol or Daltons, Num. H-bond donors ≤5, Num. H-bond acceptors ≤10 and partition coefficient log Po/w ≤5. To improve the prediction accuracy, sometimes molar refractivity from 40 to 130 is also considered as a rule of Lipinski’s rule of five (93, 94). According to our study, withaferin A and camptothecin followed all the primary criteria of Lipinski’s rule of five, reflecting the fact that they might be the best phytochemicals to combat HIV infection among the selected ligands. On the other hand, both hypericin and theaflavin violated more than one primary criterion. Hypericin has a molecular weight of 504.44 g/mol and 06 H-bond donors. Theaflavin has a molecular weight of 564.49 g/mol, 12 H-bond acceptors, and 09 H-bond donors. Other ligands, ursolic acid and oleanolic acid, violated only one primary rule each, and for this reason, they were also found to contain the properties of active drugs. These results are also in agreement with the results generated by the positive controls. Maraviroc followed Lipinski’s rule of five with one violation; however, Plerixafor was predicted to be not very active as it violated two primary criteria of Lipinski’s rule of five. Considering the molar refractivity, although it is not a primary criterion, only withaferin A and camptothecin were found to have their values within the acceptable range of 40 to 130. For this reason, among the ligands on which the druglikeness property analysis was conducted, withaferin A and camptothecin can be considered the best ligands or phytochemicals for treating HIV infections. The results of the druglikeness property analysis are listed in Table 3.

**Table 3.** Drug-likeness properties of selected compounds.

| **Drug-likeness properties** | **Rules** | **Ursolic acid** | **Oleanolic acid** | **Withaferin A** | **Hypericin** | **Camptothecin** | **Theaflavin** | **Maraviroc** | **Plerixafor** |
| --- | --- | --- | --- | --- | --- | --- | --- | --- | --- |
| **Lipinski's rule of five** | - | Yes | Yes | Yes | No | Yes | No | Yes | No |
| **Molecular weight (g/mol)** | <500 | 456.70 | 456.70 | 470.60 | 504.44 | 348.35 | 564.49 | 513.67 | 502.78 |
| **Log Po/w** | ≤5 | 5.88 | 6.06 | 3.45 | 4.26 | 2.20 | 1.31 | 4.81 | 0.68 |
| **Num. H-bond acceptors** | ≤10 | 03 | 03 | 06 | 08 | 05 | 12 | 06 | 08 |
| **Num. H-bond donors** | ≤5 | 02 | 02 | 02 | 06 | 01 | 09 | 01 | 06 |
| **Molar refractivity** | 40 - 130 | 136.91 | 136.65 | 127.49 | 144.83 | 95.31 | 143.98 | 145.84 | 182.21 |

### 3.3 ADME/T Analysis

ADMET virtual test was conducted in this study to predict the absorption, distribution, metabolism, excretion, and toxicity of the best selected ligands or phytochemicals (Table 4). The Caco-2 permeability represents the permeability and absorption of a chemical compound through the intestinal cell barrier (95). The study found that camptothecin and maraviroc, a positive control, showed predicted Caco-2 permeability, a key efflux membrane transporter in the body, hindering therapeutic agent uptake. So, limiting the activity of p-gp proteins is desirable for the therapeutic delivery of a drug (90). The study found that withaferin A, camptothecin, theaflavin, and maraviroc inhibited p-gp protein, with camptothecin being the p-gp substrate. Other ligands and positive control faced no issues during cellular uptake. Plasma protein binding (PPB) is a parameter used in deciding the distribution fate of a drug and the lesser bound a drug is, the more effectively it can be distributed to its target organs (91). Camptothecin exhibits undesirable plasma protein binding, while ligands and controls are BBB permeable. Cytochrome P450 enzymes are crucial for drug metabolization, with CYP450 substrates more effectively metabolized. Furthermore, CYP450 inhibition can have beneficial or harmful outcomes depending on the pharmacokinetic properties of the specific drug molecule (92). The study found that withaferin A, a phytochemical, was found to be a substrate for two CYP450 enzymes, indicating that it provided better metabolic activity results than other phytochemicals. The study found that hypericin had the highest excretion half-life of 3.306 hours, followed by theaflavin. Ursolic acid and oleanolic acid were predicted to be non-hERG blockers, hepatotoxic, non-Ames mutagenic, and non-toxic to the liver. Camptothecin and theaflavin were found to be toxic in three parameters. Considering all the parameters of ADMET analysis, withaferin A could be declared as the best-performing phytochemical, followed by ursolic acid and oleanolic acid.

**Table 4.**
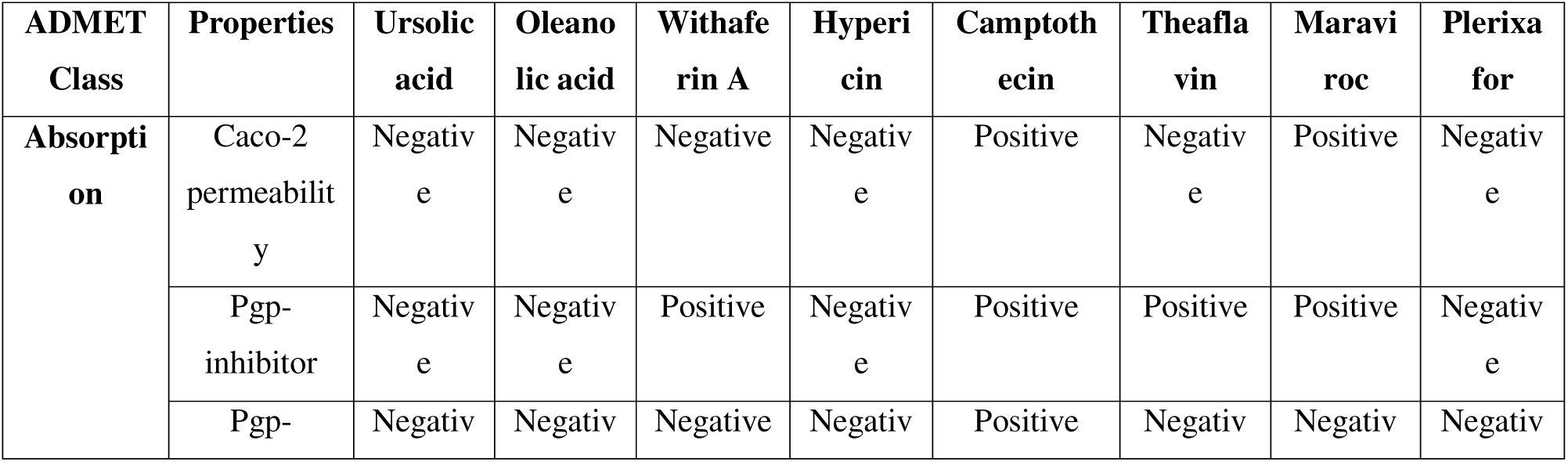
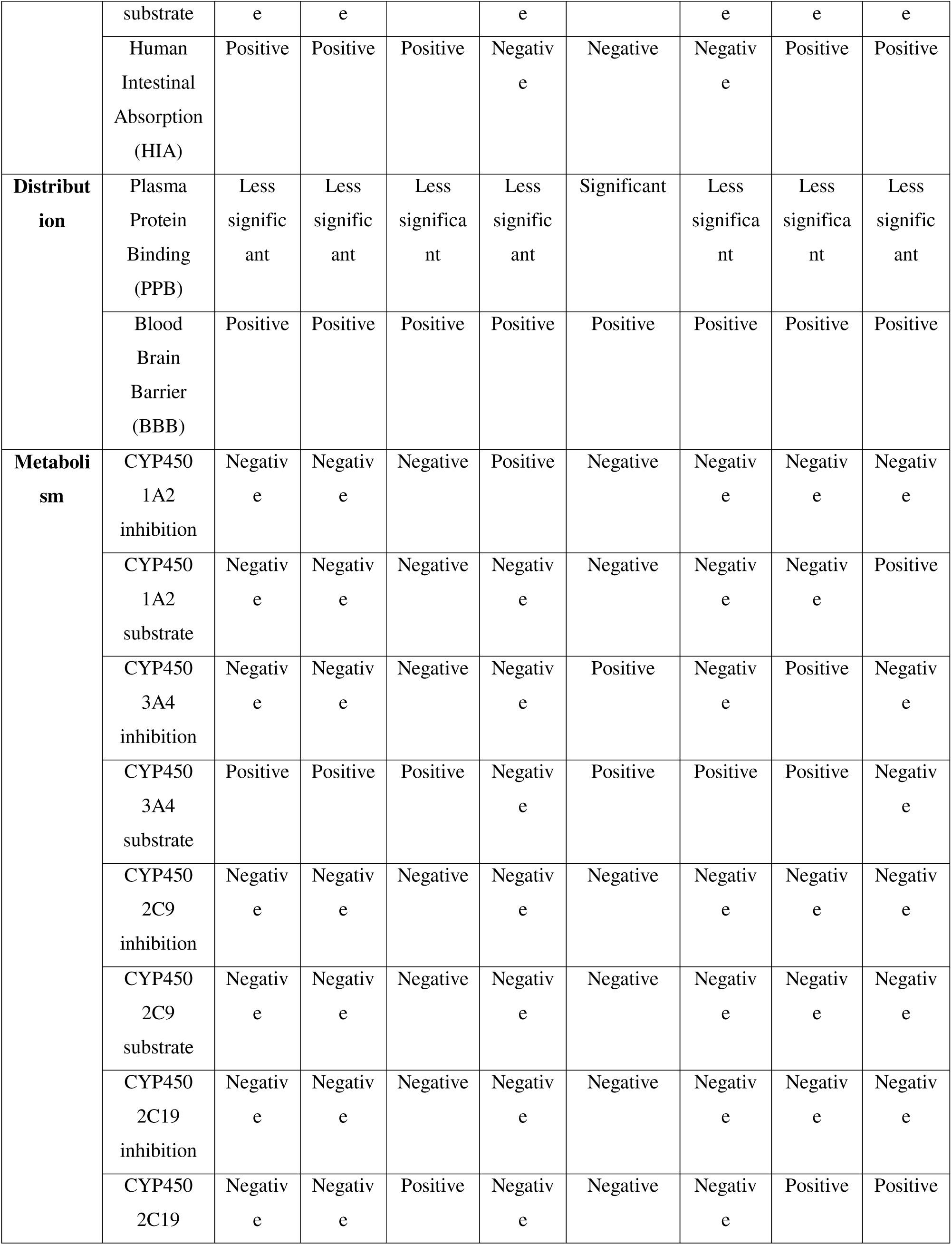
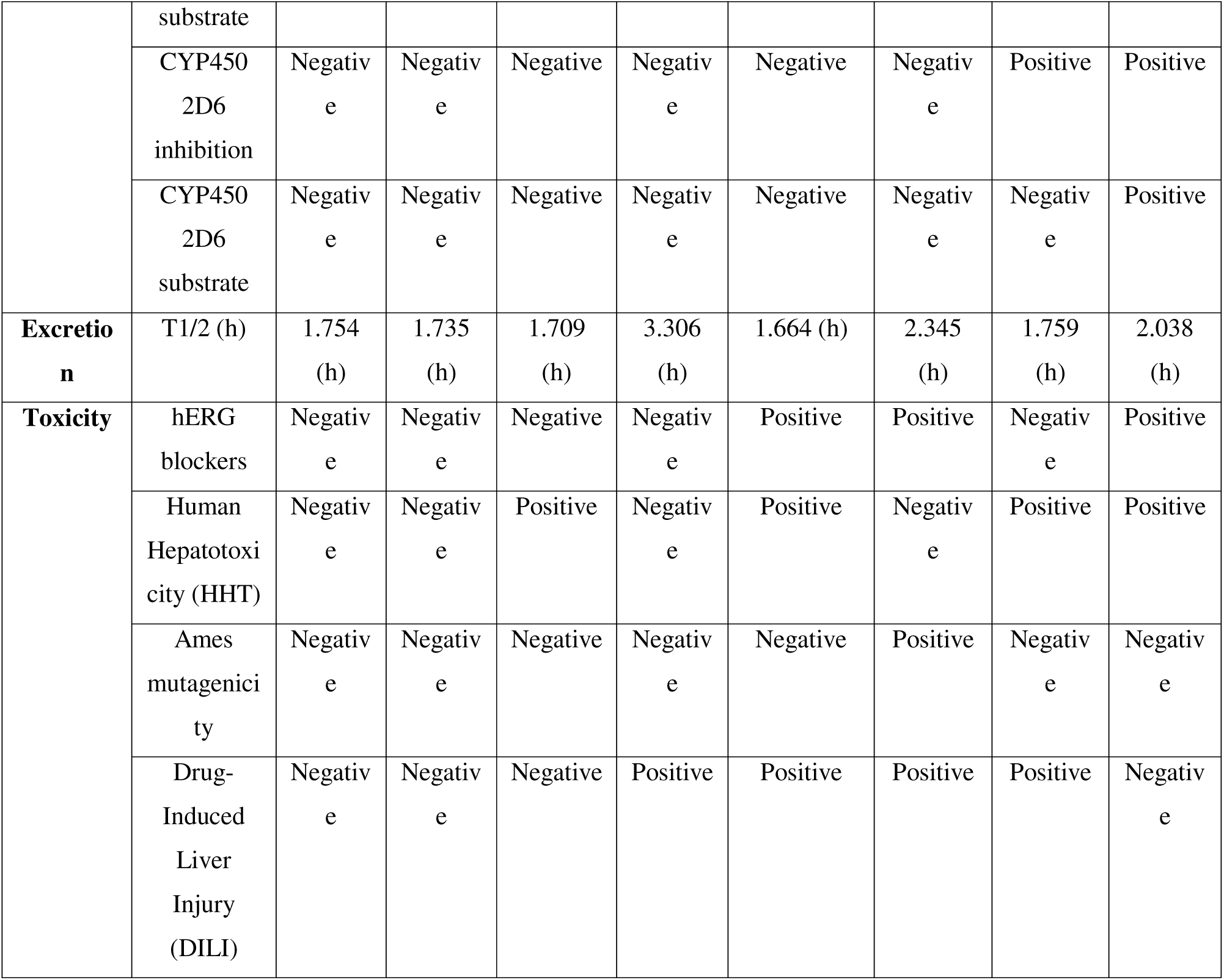
ADMET Analysis.

### 3.4 Pass Prediction

PASS forecast is a technique for investigating the natural exercises of drug-like molecules by considering two likelihood factors, i.e., Dad addressing the likelihood of a compound being dynamic and Pi addressing the possibility of a compound being dormant. Any Pa value more noteworthy than 0.7 portrays a higher probability of a compound showing a specific action or trademark. Hence, the Pa>0.7 edge gives a profoundly solid movement forecast (82). The results of the PASS prediction analysis of this study are listed in Table 5.

**Table 5.**
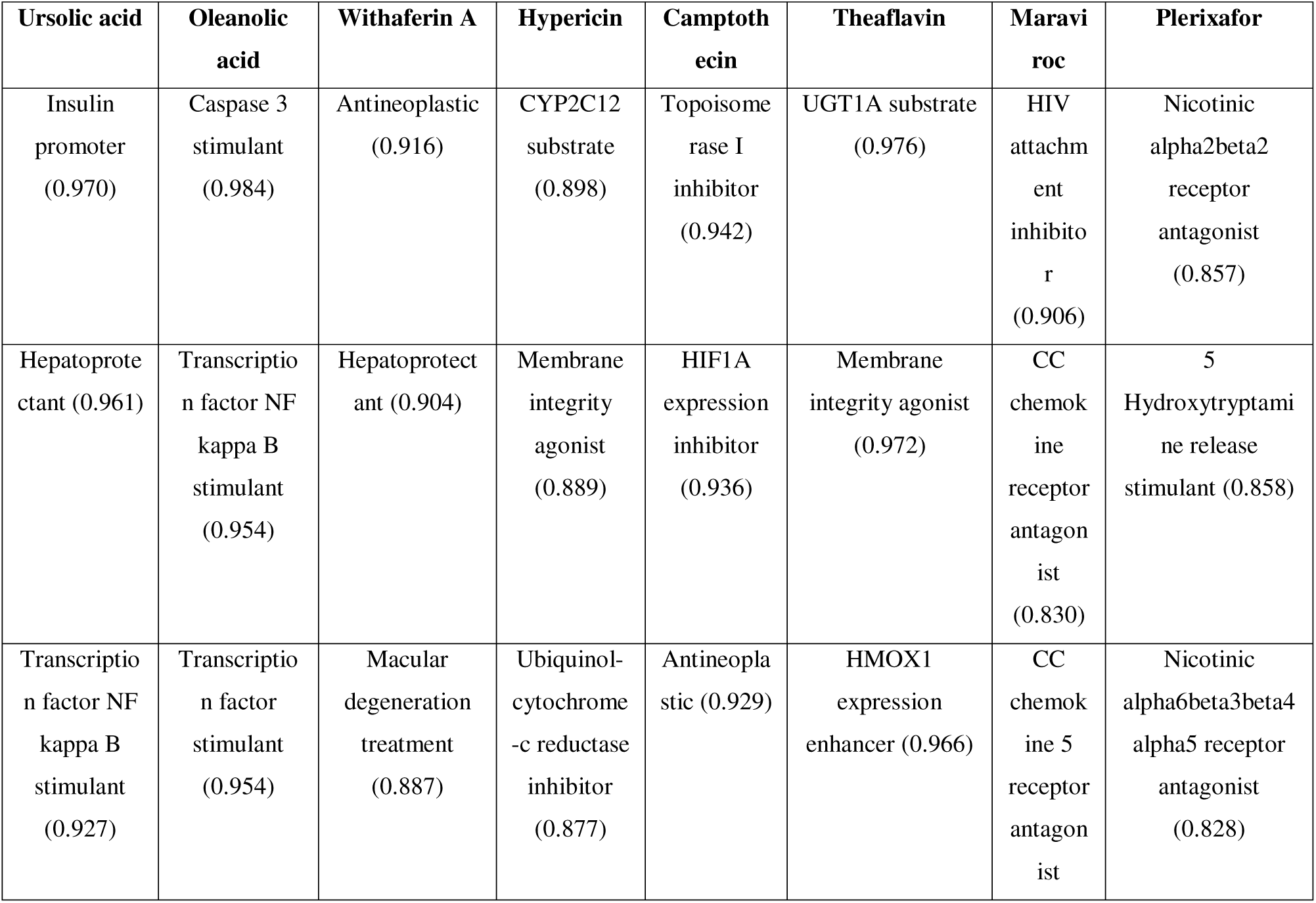
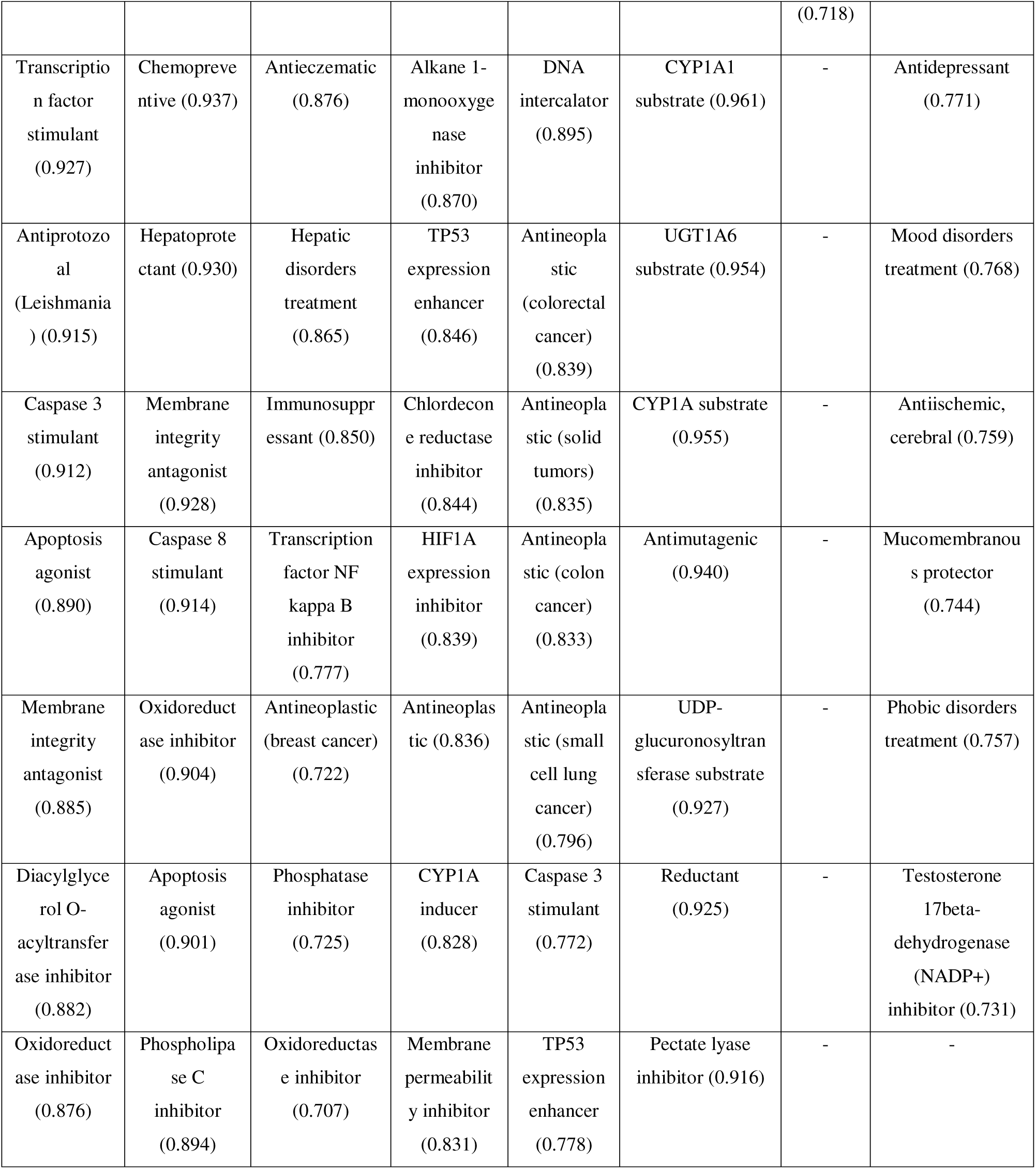
Pass Prediction Analysis.

This research showed that apart from potential anti-HIV properties (based on the molecular docking study), ursolic acid, oleanolic acid, and withaferin A might have hepatoprotective activity. We can assume that at a certain dosage, these phytochemicals can exert their protective activities on the liver as phytochemicals are proven to have hepatoprotective activities at a certain dosage (96). The phytochemicals were also found to have antitumor and anticancer activities. For example, ursolic acid, oleanolic acid, and camptothecin were predicted to have the capability to act as Caspase-3 and Caspase-8 stimulants. Caspases mediate programmed cell death or apoptosis and thus depending on the cellular context, they are antitumorigenic in nature. Their activity is used as a marker for measuring the efficacy of cancer therapy (97). Again, hypericin and camptothecin were predicted to inhibit the expression of HIF1A and enhance the expression of TP53. Studies have found that the HIF1A expression increases in a wide variety of cancers in humans (98, 99). TP53 is a tumor suppressor and it is the most commonly mutated gene in many human cancers (100). Camptothecin can be declared as the best phytochemical according to the PASS prediction analysis as it was predicted to exhibit antineoplastic activities against multiple human cancers and found to possess HIF1A expression inhibitory, TP53 expression enhancing, and Caspase 3 stimulating properties.

### 3.5 P450 SOM Prediction

The P450 site of metabolism (SOM) analysis was conducted to find out the sites of the selected phytochemicals that are more likely to be metabolized by CYP450 3A4, CYP450 2D6, and CYP450 2C9 (Table 6). Emphasis was mainly given on the lowest activation energy and lowest score because lower (activation) energy and lower score represent the higher probability of a site of a molecule being metabolized. Therefore, the sites and the molecules having lower energies and lower scores tend to be better metabolized by the CYP450 enzymes (101, 102). The P450 SOM analysis generated results that are very much in agreement with the metabolism part of the previously done ADMET analysis. According to the SOM analysis, withaferin A showed the best results with its predicted lowest energy of 52.4 and the lowest score of 43.4 CYP450 3A4 section. Therefore, withaferin A should be better metabolized by the CYP450 3A4 enzyme among the selected phytochemicals. In the ADMET analysis, withaferin A also generated the best results among the selected phytochemicals. Furthermore, according to the ADMET analysis, hypericin was the only phytochemical that might not be metabolized well by the CYP450 3A4 enzyme. According to the SOM analysis, hypericin provided the highest energy and highest score, representing its less active nature than the CYP450 3A4 enzyme. Again, according to the ADMET analysis, all the phytochemicals showed poor performance in terms of metabolizing by the enzymes compared to the positive controls. Likewise, the SOM analysis also showed that the energies and the scores generated by the phytochemicals were much higher than those of the positive controls, reflecting their unsatisfactory performance relative to the positive controls. And among the phytochemicals, with lower energies and lower scores in terms of all the three CYP450 enzymes, withaferin A could be declared as the best performing agent.

**Table 6.**
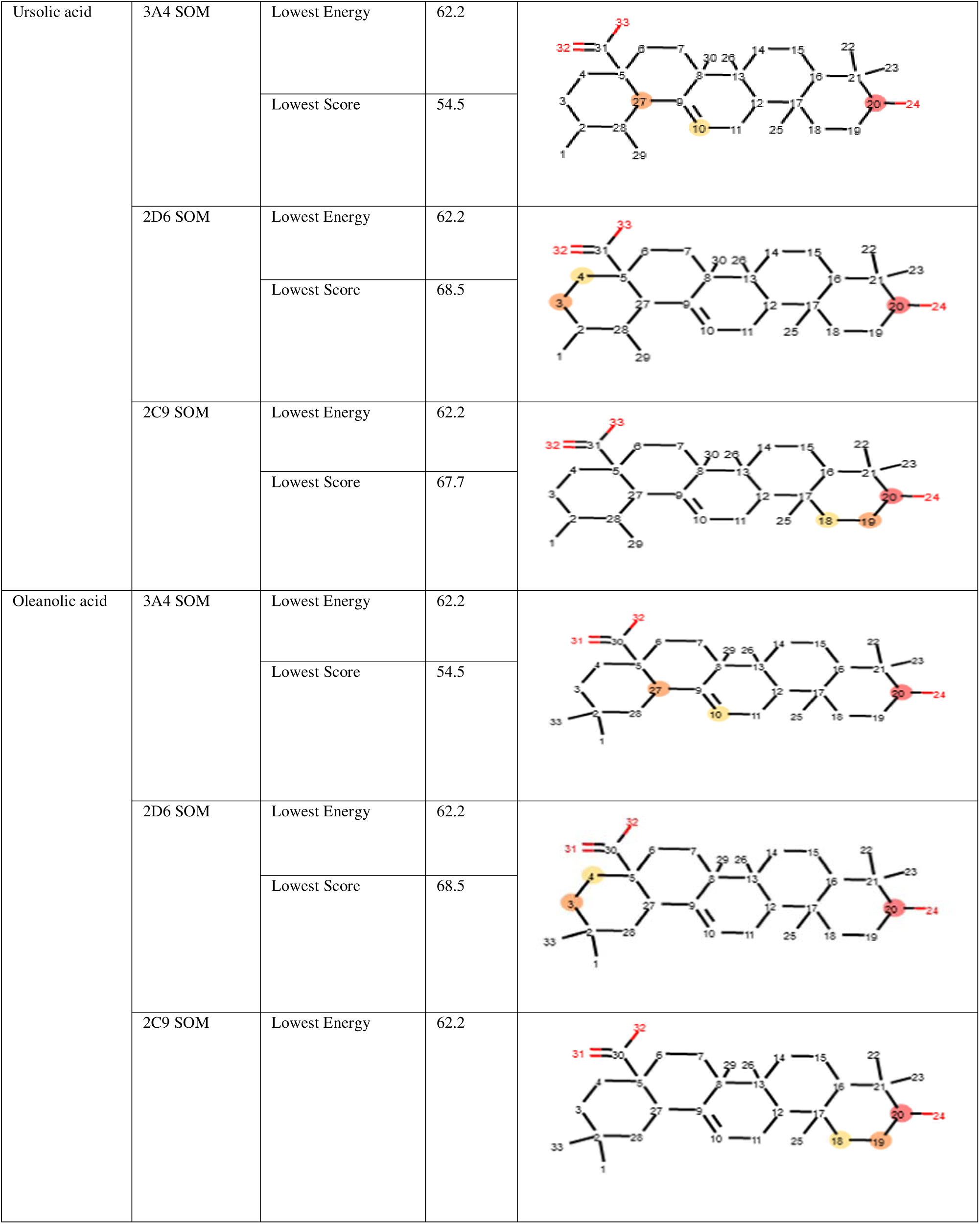
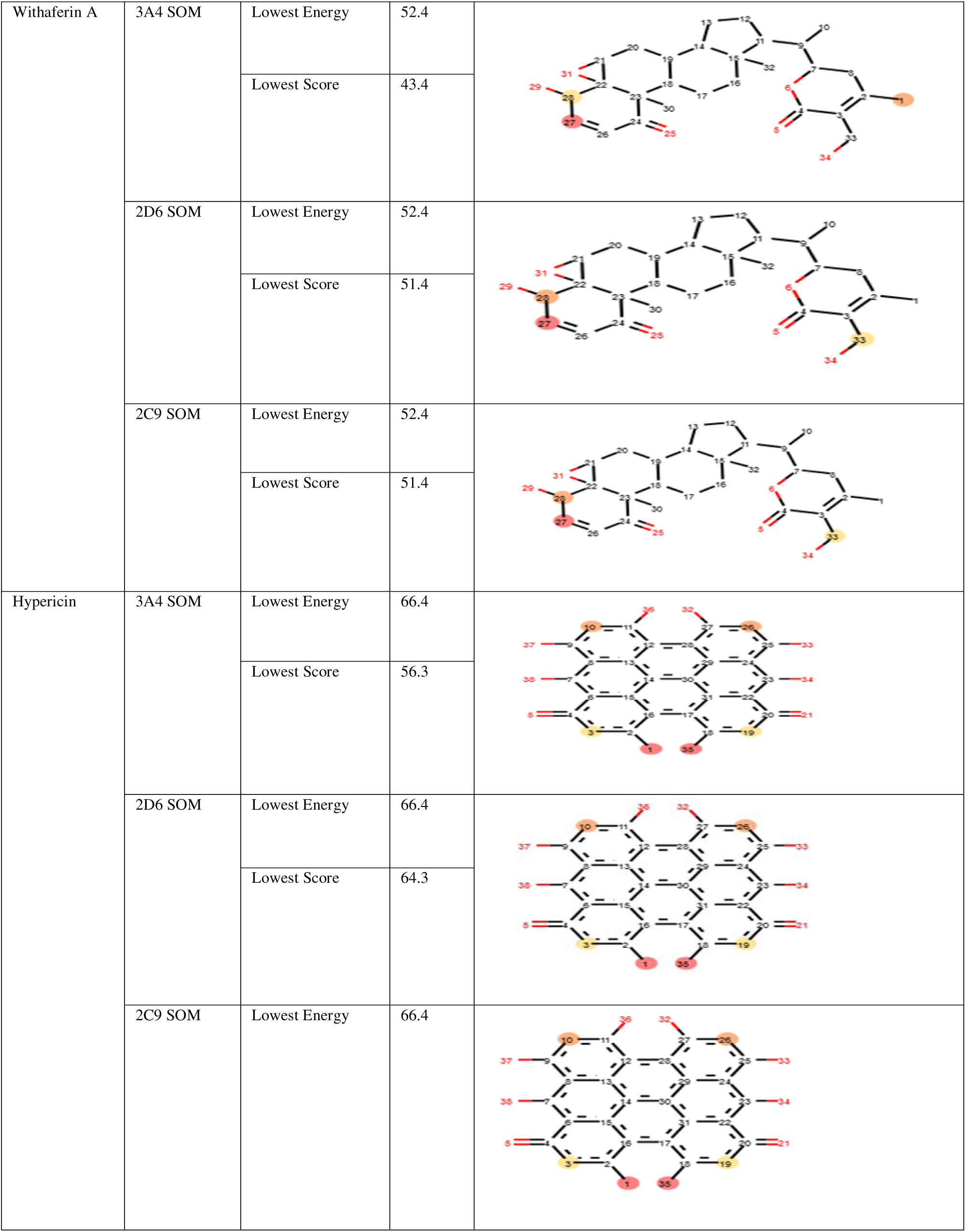
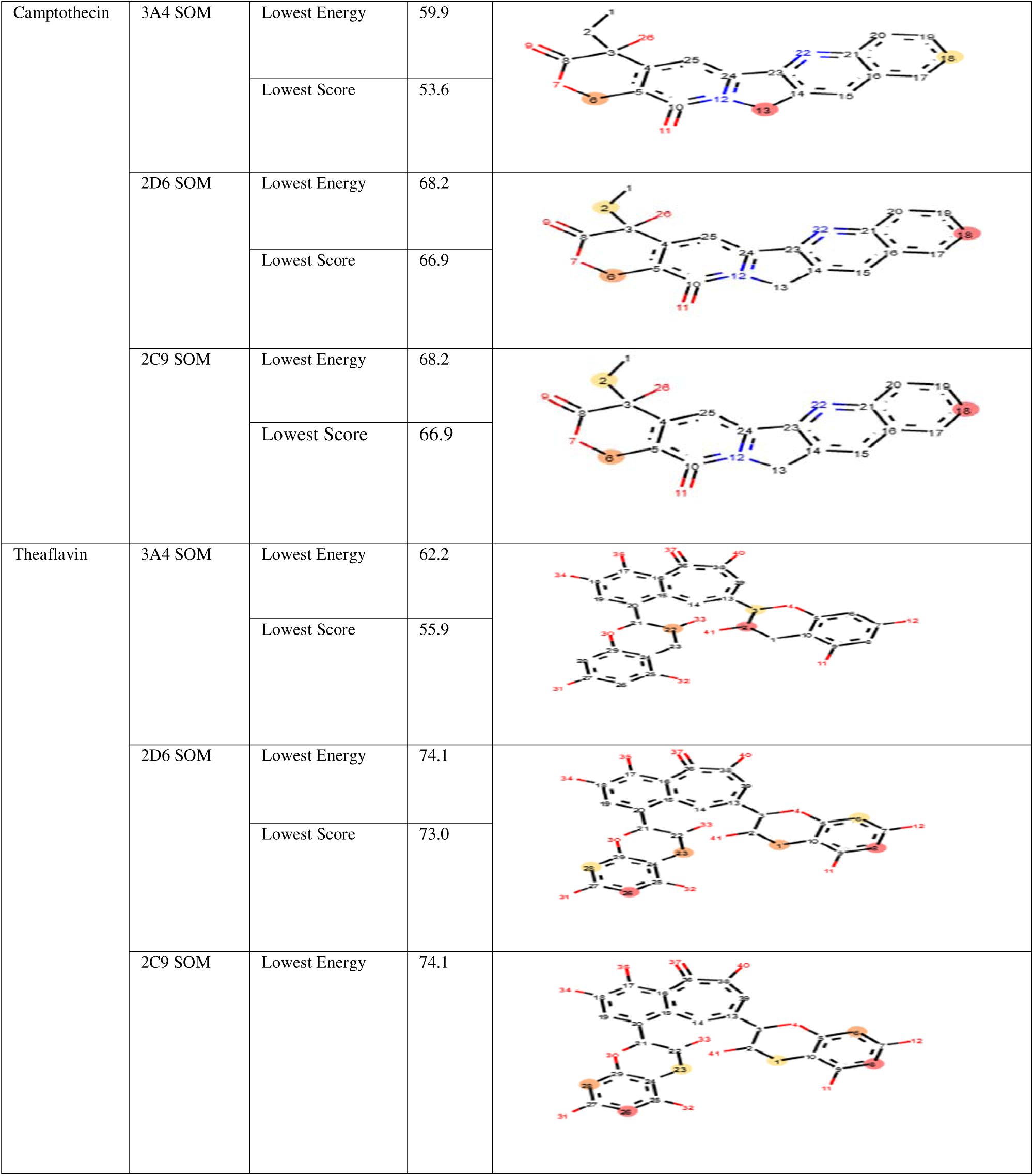
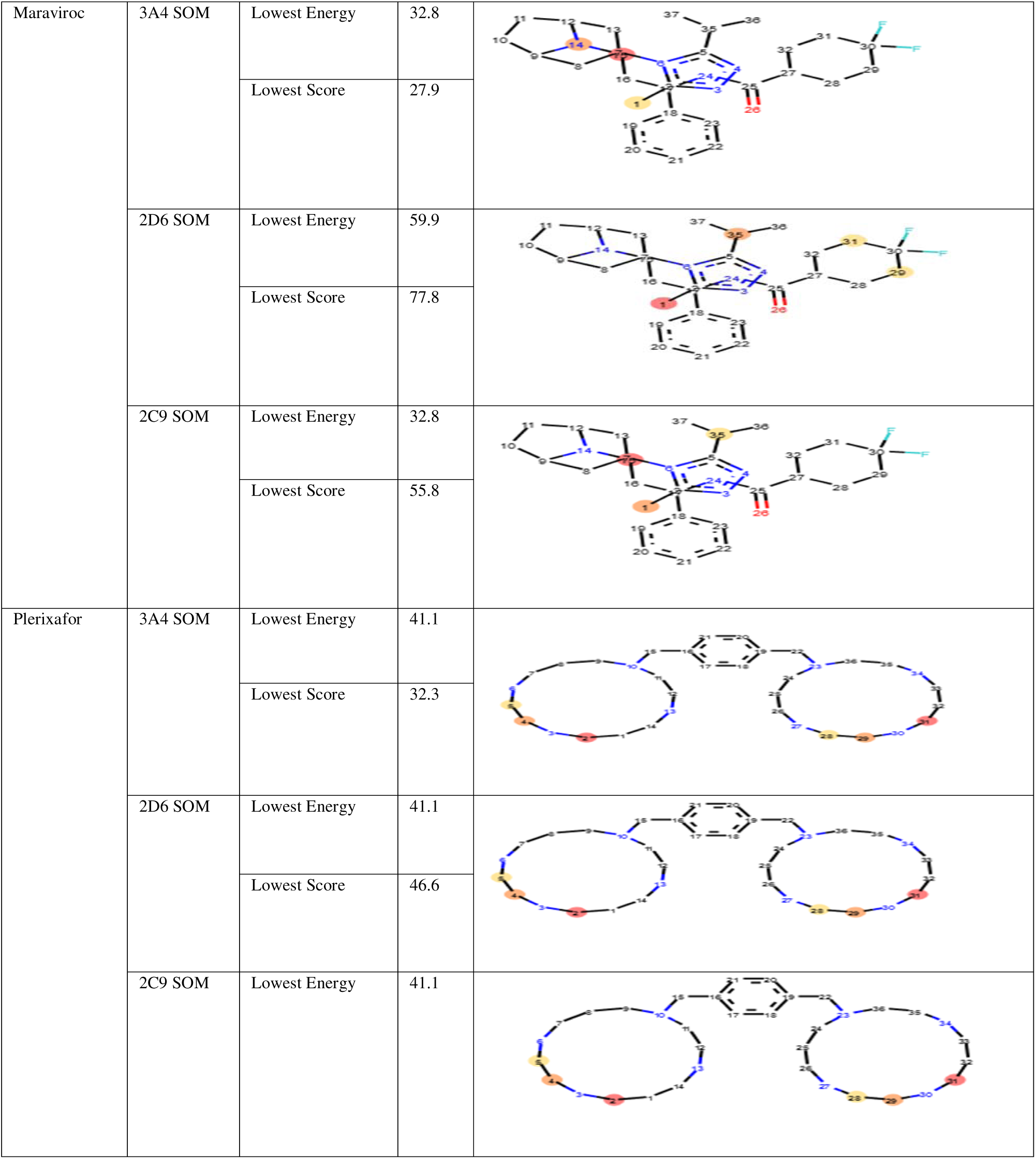
P450 SOM Prediction.

### 3.6 Molecular Dynamic Simulation

XmGrace package was used to analyze the MD trajectory parameters (103). During the MD run, the RMSD is determined between a characterized beginning stage of the recreation and every single succeeding edge (104).

The analysis of various compounds (Table 7) revealed that the average RMSD value for various compounds was 0.2287 nm for the 3ODU complex, 0.0453 nm for 3ODU-Hypericin, 0.0976 nm for 3ODU-Oleanolic Acid, 0.0844 nm for 3ODU-Theaflavin, 0.2098 nm for 3ODU-Camptothecin, 0.3455 nm for 3ODU-Plerixafor, and 0.1139 nm for the 4MBS complex. The graph (Figure 3) showed the stability of various complexes, with the 3ODU complex exhibiting the highest stability at 15 to 32 nanoseconds. The 3ODU-Camptothecin complex had the highest stability at 24 to 50 nanoseconds, while the 4MBS complex had the lowest stability at 16 to 60 nanoseconds. The RMSD is valuable in analyzing the time-dependent motion of a given structure throughout the simulation (105). In this way, a plateau of RMSD values indicates that the structure fluctuates around a stable average conformation, which can be observed in all of the performed MD simulations.

**Table 7.** RMSD Analysis.

| Value | RMSD (nm) in C-α | | |
| --- | --- | --- | --- |
|  | Average (nm) | Minimum (nm) | Maximum (nm) |
| 3ODU | 0.2287 | 0.1121 | 0.3453 |
| 3ODU-Hypericin | 0.0453 | 0.0234 | 0.6112 |
| 3ODU-Oleanolic Acid | 0.0976 | 0.0532 | 0.1421 |
| 3ODU-Ursolic Acid | 0.0844 | 0.0613 | 0.1075 |
| 3ODU-Theaflavine | 0.2098 | 0.1027 | 0.3169 |
| 3ODU-Camptothecin | 0.3455 | 0.3012 | 0.3898 |
| 3ODU-Plerixafor | 0.1139 | 0.0241 | 0.2037 |
| 4MBS | 0.1943 | 0.1165 | 0.2721 |
| 4MBS-Withaferin A | 0.1765 | 0.0642 | 0.2888 |
| 4MBS-Oleanolic Acid | 0.1108 | 0.0452 | 0.1742 |
| 4MBS-Maraviroc | 0.3436 | 0.1129 | 0.5743 |
| 4MBS-Ursolic Acid | 0.0887 | 0.0312 | 0.1462 |
| 4MBS-Hypericin | 0.0976 | 0.0431 | 0.1521 |

The local flexibility in light of buildup relocations during the MD simulation can be characterized by utilizing RMSF values (103). The RMSF value (Table 8) stability for 3ODU and 4MBS complexes with the chosen molecules is shown in the graph (Figure 4). These complexes fall within the following stability ranges: 800–900, 3500–3700, 1000–2700, 1700–2200, 1000–2000, 2000–3000, and 1500–3000. The average RMSF value for various compounds was found to be 0.2813 nm for 3ODU-Hypericin, 0.1721 nm for 3ODU-Oleanolic Acid, 0.3895 nm for 3ODU-Ursolic Acid, and 0.3237 nm for 3ODU-Theaflavin. The 4MBS-Hypericin complex had an average of 0.2813 nm, while 3ODU-Camptothecin had an average of 0.4246 nm. Other compounds had averages of 0.3421 nm, 0.3421 nm, and 0.3782 nm. The average RMSF value is higher in the 3ODU-Camptothecin complex (0.4246 nm) and lowest in the 4MBS-Oleanolic Acid complex (0.1654 nm) Table 8. Higher RMSF values represent more flexible movements, whereas lower RMSF values represent movements that are more constrained in regard to average locations during simulation (105).

**Figure 4:**
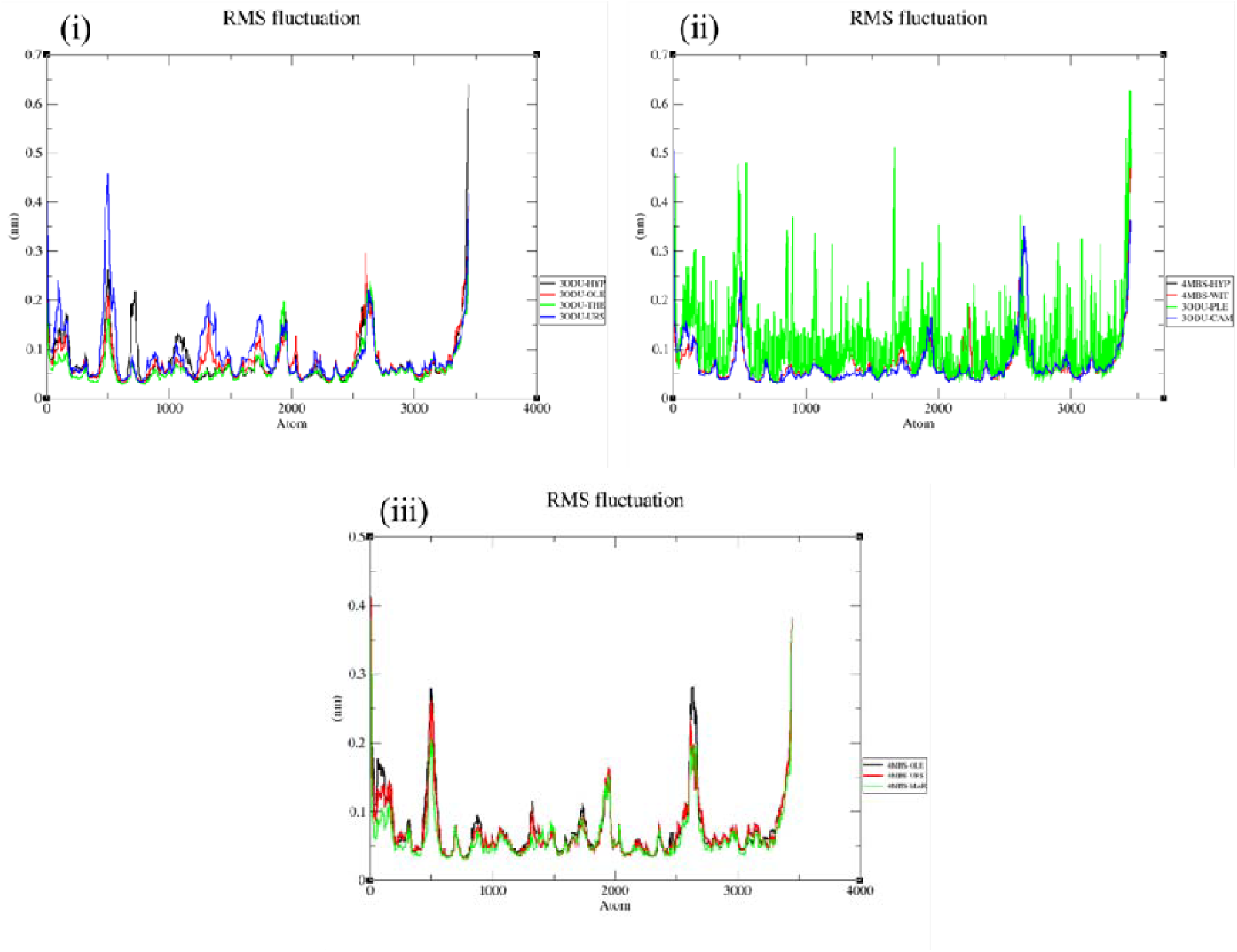
RMSF (Root Mean Square Fluctuation) of (i) 3ODU-Hypericin (black), 3ODU-Oleanolic Acid (Red), 3ODU-Theaflavin (Green), 3ODU-Ursolic Acid (Blue) (ii) 4MBS-Hypericin (Black), 4MBS-Withaferin A (Red), 3ODU-Plerixafor (Green), 3ODU-Camptothecin (Blue) (iii) 4MBS-Oleanolic Acid (Black color),4MBS-Ursolic Acid (Red), 4MBS-Maraviroc (Green)

**Table 8.** RMSF Analysis.

| Value | RMSF (nm) in C-α | | |
| --- | --- | --- | --- |
|  | Average (nm) | Minimum (nm) | Maximum (nm) |
| 3ODU-Hypericin | 0.2813 | 0.1121 | 0.4505 |
| 3ODU-Oleanolic Acid | 0.1721 | 0.1232 | 0.2211 |
| 3ODU-Ursolic Acid | 0.3895 | 0.0523 | 0.7267 |
| 3ODU-Theaflavine | 0.3237 | 0.0927 | 0.5547 |
| 3ODU-Camptothecin | 0.4246 | 0.2012 | 0.6480 |
| 3ODU-Plerixafor | 0.3421 | 0.1241 | 0.5601 |
| 4MBS-Withaferin A | 0.3782 | 0.0322 | 0.7242 |
| 4MBS-Oleanolic Acid | 0.1654 | 0.0452 | 0.2856 |
| 4MBS-Maraviroc | 0.2836 | 0.0424 | 0.5248 |
| 4MBS-Ursolic Acid | 0.1975 | 0.0922 | 0.3028 |
| 4MBS-Hypericin | 0.2813 | 0.1431 | 0.4229 |

The radius of gyration (RG) is commonly described as the root mean square distance of a group of atoms from their shared center of mass, with the added consideration of mass weighting (106). The stability of the radius of the gyration value for the complexes is displayed on the graph (Figure. 5). 3ODU-Hypericin has the longest stability, ranging from 20 to 50 nanoseconds. Shorter stability times of 40–80, 30–60, and 50–70 nanoseconds are seen in other compounds. These compounds have an average Rg value of 1.4321 nm. The Rg value of the 4MBS-Hypericin combination is greater at 2.4813 nm. As a result, the RG score was higher in the 4MBS-Hypericin complex (2.4813 nm) and lower in the 3ODU-Hypericin complex (1.4321 nm) (Table 9). The findings suggest that the protein-ligand complex exhibited lower compactness compared to the apoprotein.

**Figure 5:**
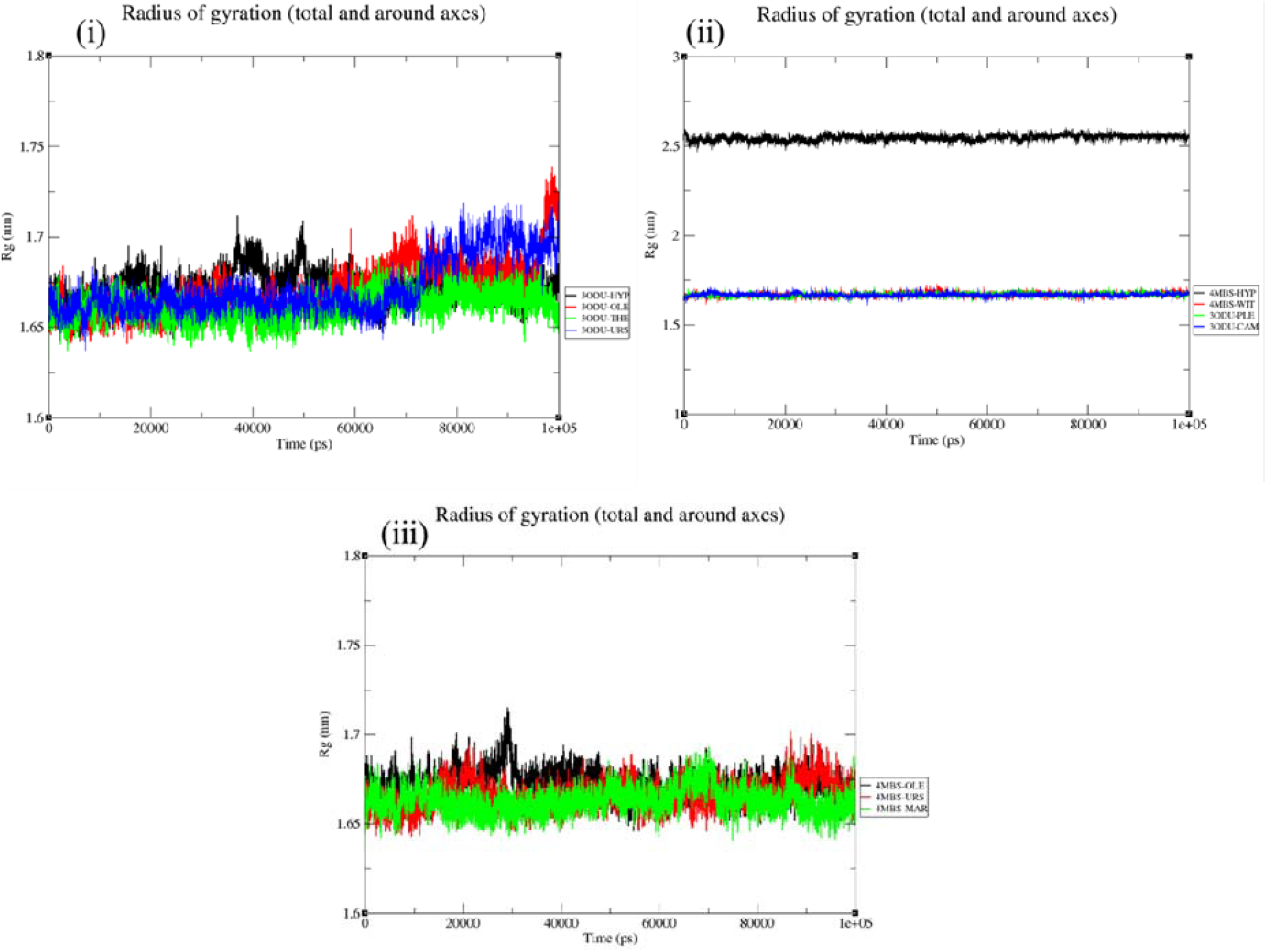
Rg (Radius of Gyration) of (i) 3ODU-Hypericin (black), 3ODU-Oleanolic Acid (Red), 3ODU-Theaflavin (Green), 3ODU-Ursolic Acid (Blue) (ii) 4MBS-Hypericin (Black), 4MBS-Withaferin A (Red), 3ODU-Plerixafor (Green), 3ODU-Camptothecin (Blue) (iii) 4MBS-Oleanolic Acid (Black color),4MBS-Ursolic Acid (Red), 4MBS-Maraviroc (Green) (Brown)

**Table 9.** Radius of Gyration Analysis.

| Value | Radius of Gyration (nm) |  |  |
| --- | --- | --- | --- |
|  | Average (nm) | Minimum (nm) | Maximum (nm) |
| 3ODU-Oleanolic Acid | 1.5721 | 1.4232 | 1.7211 |
| 3ODU-Hypericin | 1.4321 | 1.2111 | 1.6721 |
| 3ODU-Ursolic Acid | 1.6895 | 1.4523 | 1.9258 |
| 3ODU-Theaflavine | 1.6637 | 1.3227 | 2.0047 |
| 3ODU-Camptothecin | 1.6246 | 1.4012 | 1.8480 |
| 3ODU-Plerixafor | 1.5421 | 1.4241 | 1.6601 |
| 4MBS-Withaferin A | 1.5732 | 1.4822 | 1.6642 |
| 4MBS-Oleanolic Acid | 1.6624 | 1.5452 | 1.7832 |
| 4MBS-Maraviroc | 1.7826 | 1.6414 | 1.9238 |
| 4MBS-Ursolic Acid | 1.7775 | 1.5522 | 2.0028 |
| 4MBS-Hypericin | 2.4813 | 2.4931 | 2.4695 |

SASA analysis methodology measures the solvent-accessible surface area of a protein, providing insights into its molecular interaction ability (106). Figure 6 and Table 10 illustrate the most stable compounds are 3ODU-Hypericin, 4MBS-Hypericin, 3ODU-Camptothecin, 3ODU-Plerixafor, and 4MBS-Withaferin A, while the least stable complexes are 4MBS-Maraviroc and 4MBS-Oleanolic Acid. The 4MBS-Hypericin combination has the highest SASA score. The findings indicate that the apoprotein exhibits comparatively lower accessibility in comparison to the protein-ligand complex, potentially impacting their ability to engage with other molecules.

**Figure 6:**
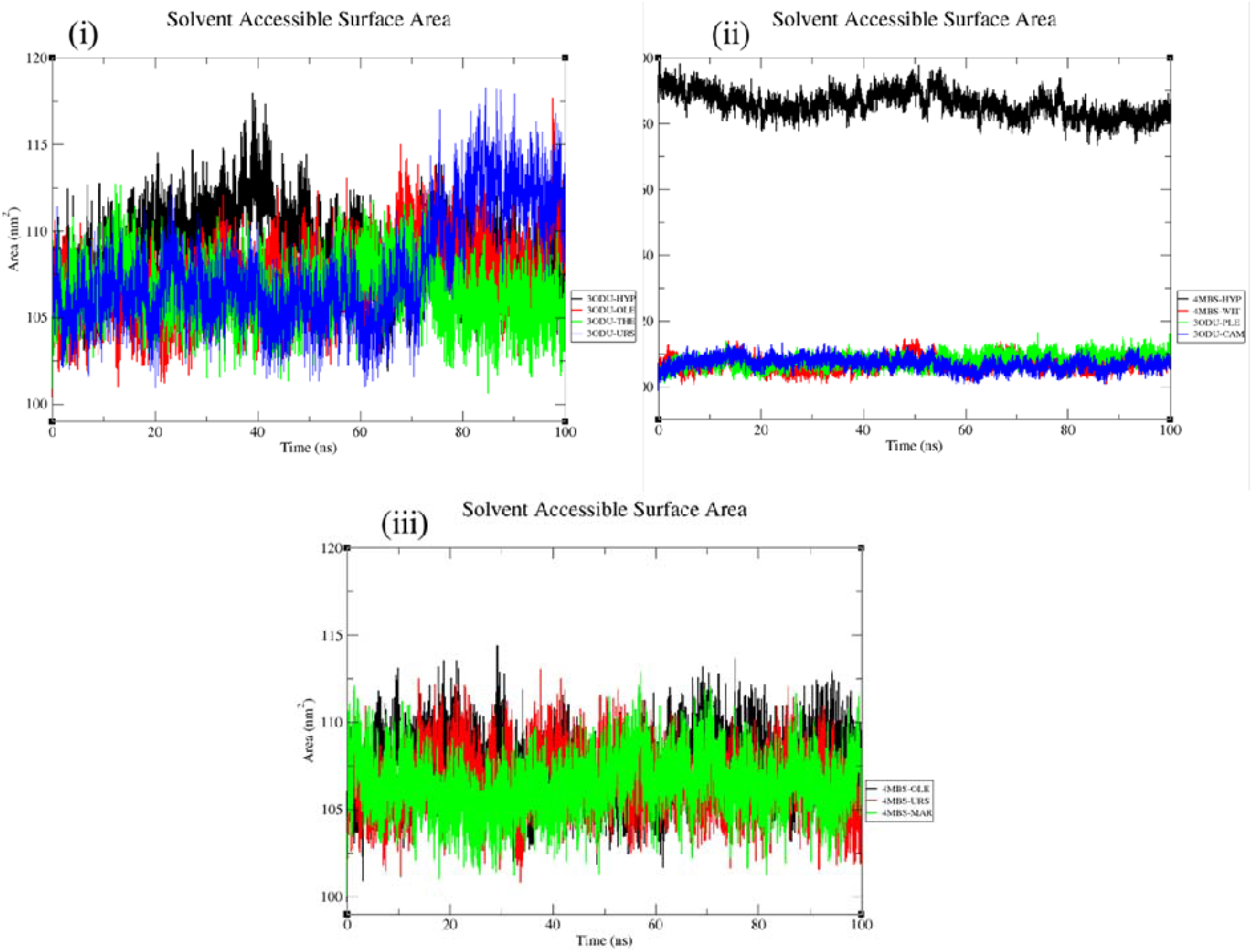
SASA (Solvent Accessible Surface Area) of (i) 3ODU-Hypericin (black), 3ODU-Oleanolic Acid (Red), 3ODU-Theaflavin (Green), 3ODU-Ursolic Acid (Blue) (ii) 4MBS-Hypericin (Black), 4MBS-Withaferin A (Red), 3ODU-Plerixafor (Green), 3ODU-Camptothecin (Blue) (iii) 4MBS-Oleanolic Acid (Black color),4MBS-Ursolic Acid (Red), 4MBS-Maraviroc (Green)

**Table 10.** SASA analysis.

| Value | Solvent Accessible Surface Area (nm<sup>2</sup>) |  |  |
| --- | --- | --- | --- |
|  | Average (nm<sup>2</sup>) | Minimum (nm<sup>2</sup>) | Maximum (nm<sup>2</sup>) |
| 3ODU-Oleanolic Acid | 111.172 | 103.123 | 119.221 |
| 3ODU-Hypericin | 108.421 | 101.2359 | 117.432 |
| 3ODU-Ursolic Acid | 110.395 | 101.053 | 119.737 |
| 3ODU-Theaflavine | 109.217 | 102.097 | 116.337 |
| 3ODU-Camptothecin | 108.426 | 105.212 | 111.644 |
| 3ODU-Plerixafor | 109.421 | 101.121 | 117.721 |
| 4MBS-Withaferin A | 106.312 | 101.032 | 111.592 |
| 4MBS-Oleanolic Acid | 111.654 | 103.042 | 120.266 |
| 4MBS-Maraviroc | 105.236 | 102.024 | 108.366 |
| 4MBS-Ursolic Acid | 105.195 | 101.022 | 109.368 |
| 4MBS-Hypericin | 179.283 | 178.131 | 181.434 |

The “Area per residue over the trajectory” is a metric used in molecular dynamics simulations and structural biology to measure the average surface area of an amino acid residue during a trajectory. It helps assess the accessibility or hiddenness of specific protein segments over time, providing insights into the dynamic behavior and interactions of biomolecules. The average “Area per residue over the trajectory” value for 3ODU-Hypericin was 0.6433 nm2, while for 3ODU-Oleanolic Acid, it was 0.4721 nm2. The analysis found that the “Area per residue over the trajectory” value is higher in the 4MBS-Ursolic Acid complex (0.8922 nm2) and lowest in the 4MBS-Withaferin A complex (0.4082 nm2) Table 11.

**Figure 7:**
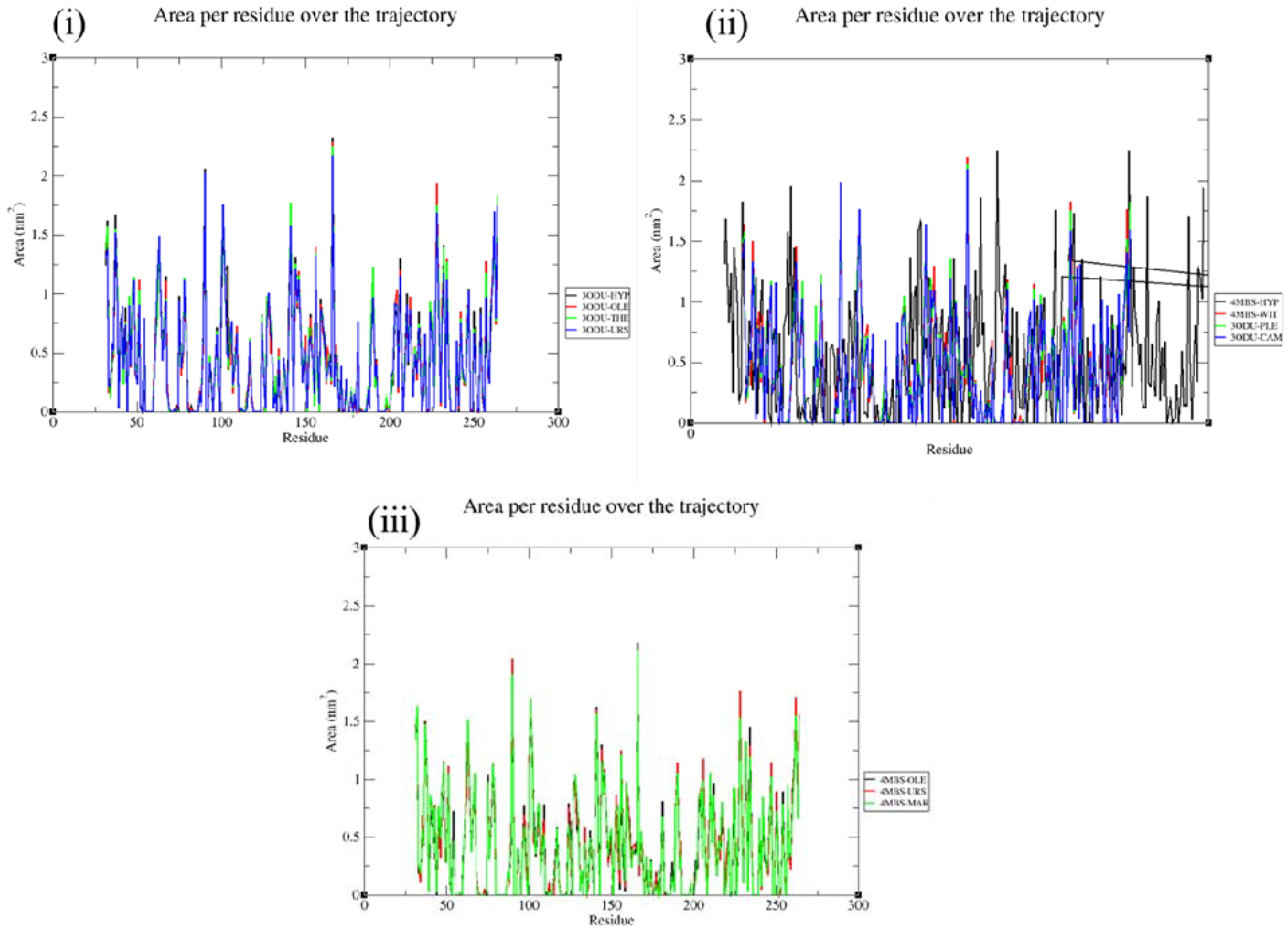
“Area per residue over the trajectory or Residue Surface Area) of (i) 3ODU-Hypericin (black), 3ODU-Oleanolic Acid (Red), 3ODU-Theaflavin (Green), 3ODU-Ursolic Acid (Blue) (ii) 4MBS-Hypericin (Black), 4MBS-Withaferin A (Red), 3ODU-Plerixafor (Green), 3ODU-Camptothecin (Blue) (iii) 4MBS-Oleanolic Acid (Black color),4MBS-Ursolic Acid (Red), 4MBS-Maraviroc (Green)

**Table 11.**
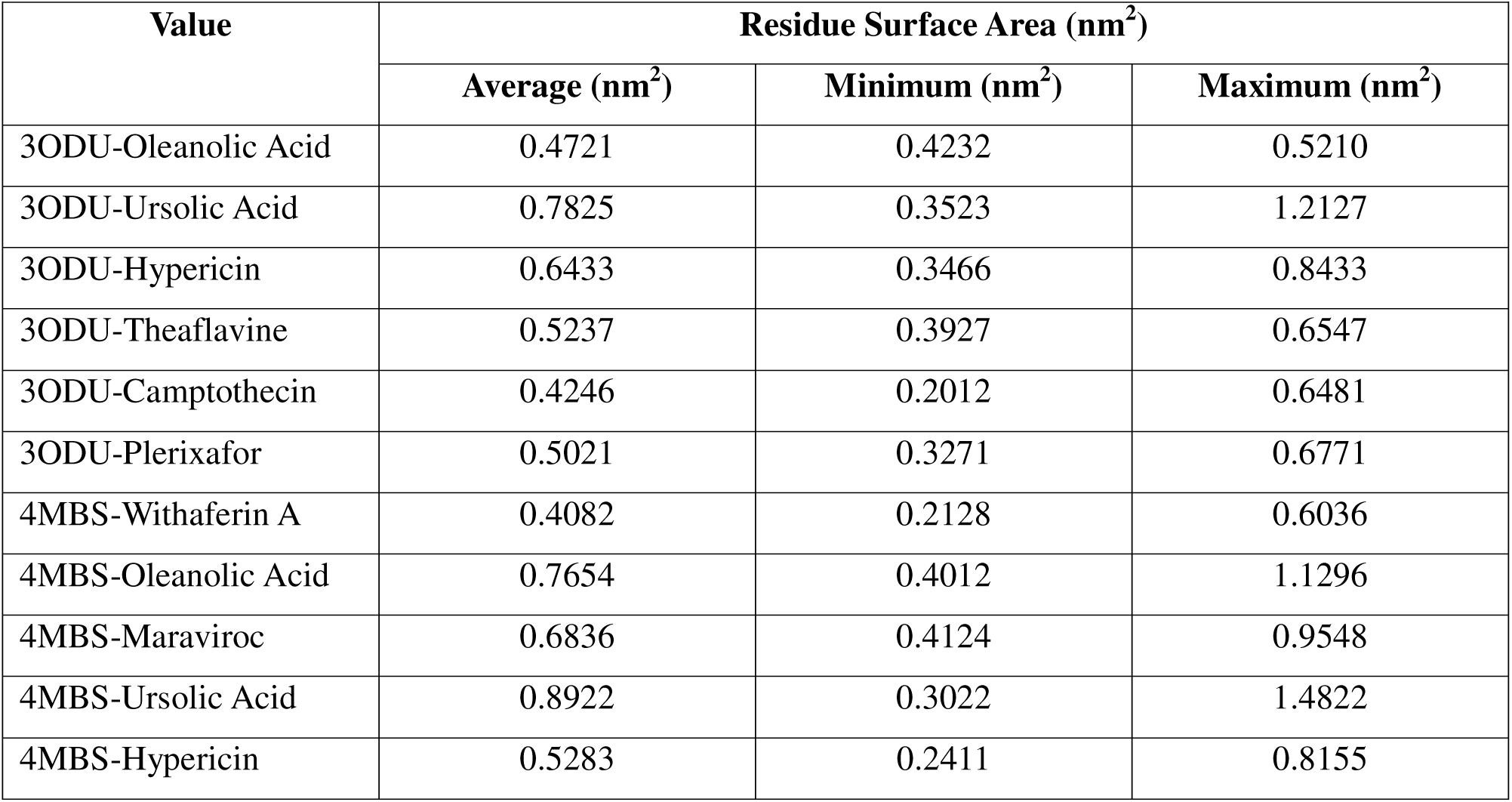
Residue Surface Area Analysis.

| Value | Residue Surface Area (nm<sup>2</sup>) |  |  |
| --- | --- | --- | --- |
|  | Average (nm<sup>2</sup>) | Minimum (nm<sup>2</sup>) | Maximum (nm<sup>2</sup>) |
| 3ODU-Oleanolic Acid | 0.4721 | 0.4232 | 0.5210 |
| 3ODU-Ursolic Acid | 0.7825 | 0.3523 | 1.2127 |
| 3ODU-Hypericin | 0.6433 | 0.3466 | 0.8433 |
| 3ODU-Theaflavine | 0.5237 | 0.3927 | 0.6547 |
| 3ODU-Camptothecin | 0.4246 | 0.2012 | 0.6481 |
| 3ODU-Plerixafor | 0.5021 | 0.3271 | 0.6771 |
| 4MBS-Withaferin A | 0.4082 | 0.2128 | 0.6036 |
| 4MBS-Oleanolic Acid | 0.7654 | 0.4012 | 1.1296 |
| 4MBS-Maraviroc | 0.6836 | 0.4124 | 0.9548 |
| 4MBS-Ursolic Acid | 0.8922 | 0.3022 | 1.4822 |
| 4MBS-Hypericin | 0.5283 | 0.2411 | 0.8155 |

The study of hydrogen bonds can direct the alteration of a lead molecule to increase its activity and give valuable insights into the stability of a ligand-protein complex. Figure 8 shows the stability of the SASA value for various complexes, including 3ODU-Hypericin, 3ODU-Oleanolic Acid, 3ODU-Ursolic Acid, and 3ODU-Theaflavin. The stability ranges from 10 to 40 nanoseconds for 3ODU-Camptothecin, 30 to 40 nanoseconds for 3ODU-Plerixafor, 55 to 75 nanoseconds for 4MBS-Hypericin, 20 to 50 nanoseconds for 4MBS-Withaferin A, 50 to 80 nanoseconds for 4MBS-Oleanolic Acid, and 30 to 60 nanoseconds for 4MBS-Ursolic Acid.

**Figure 8:**
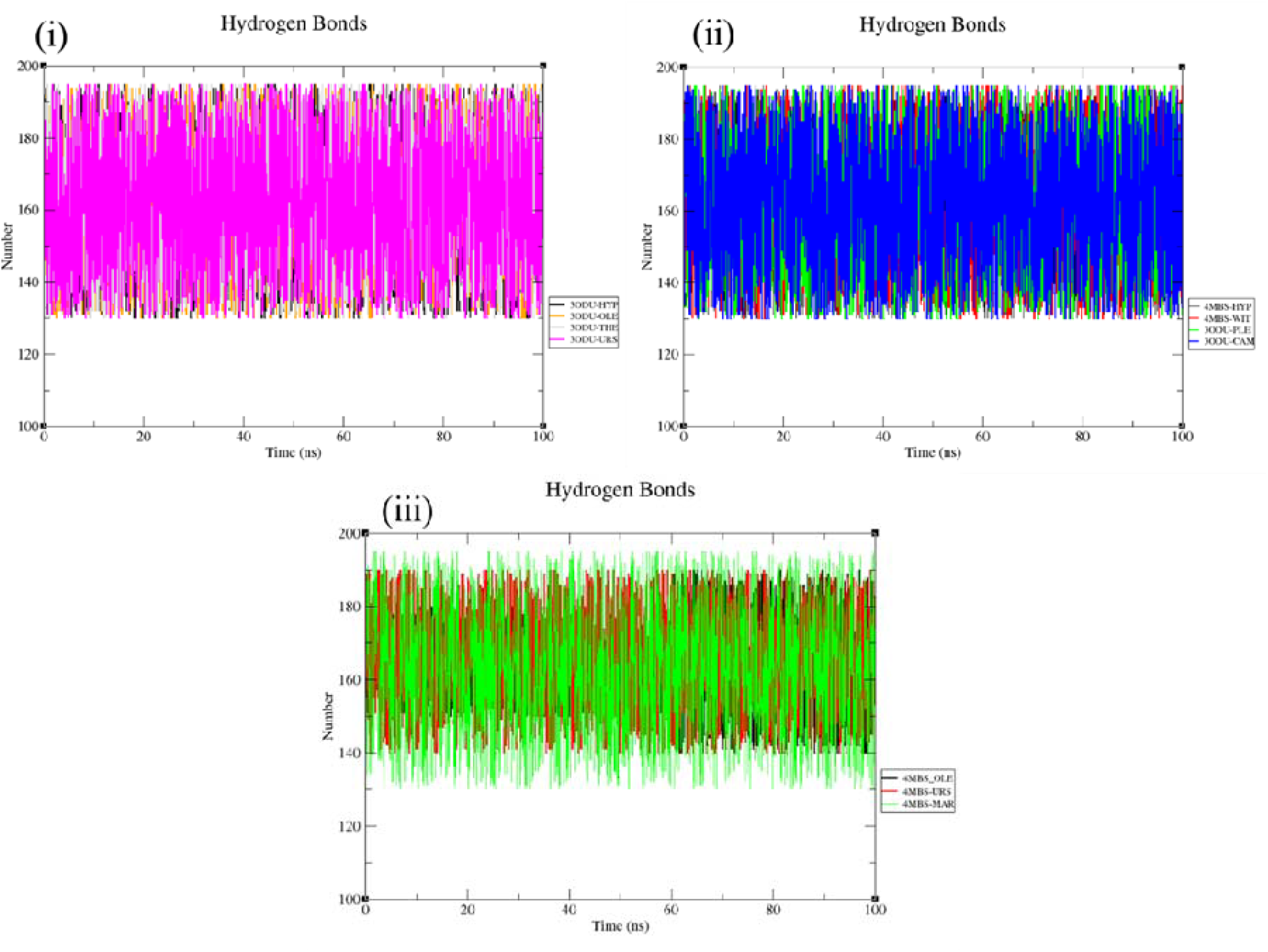
Hydrogen Bonds of (i) 3ODU-Hypericin (black), 3ODU-Oleanolic Acid (Orange), 3ODU-Theaflavin (Grey), 3ODU-Ursolic Acid (Pink) (ii) 4MBS-Hypericin (Black), 4MBS-Withaferin A (Red), 3ODU-Plerixafor (Green), 3ODU-Camptothecin (Blue) (iii) 4MBS-Oleanolic Acid (Black color),4MBS-Ursolic Acid (Red), 4MBS-Maraviroc (Cyan)

**Figure 9.**
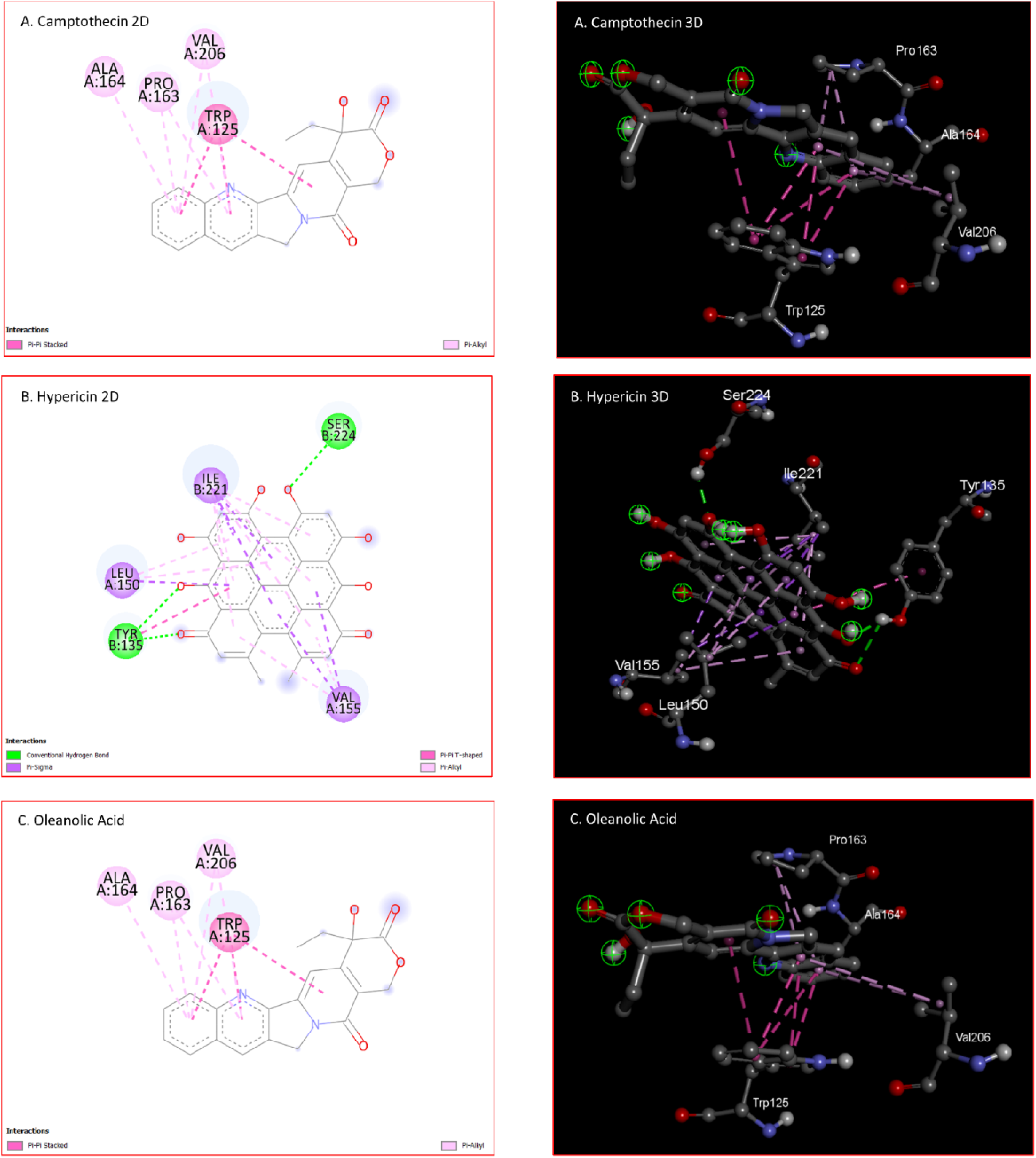
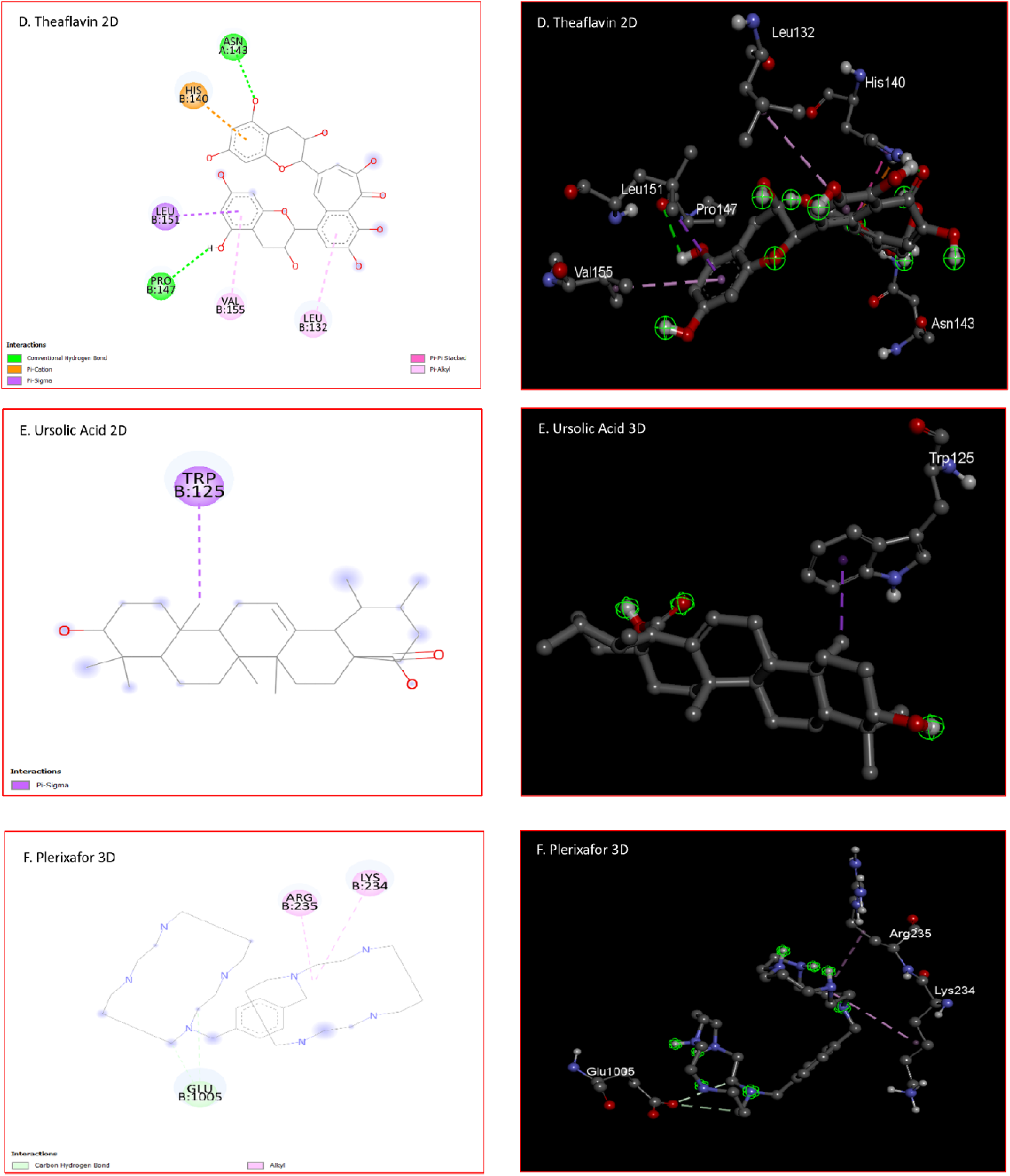
(i). 2D and 3D images show the bonds and amino acids interacting with the five best-selected ligands with the 3ODU receptor (A. Camptothecin, B. Hypericin, C. Oleanolic acid, D. Theaflavin and E. Ursolic acid, and F. Plerixafor). Interacting amino acid residues of the target molecule are outlined in the diagram, and dotted lines depict the interaction between the ligand and receptor.

**Figure 9 (ii).**
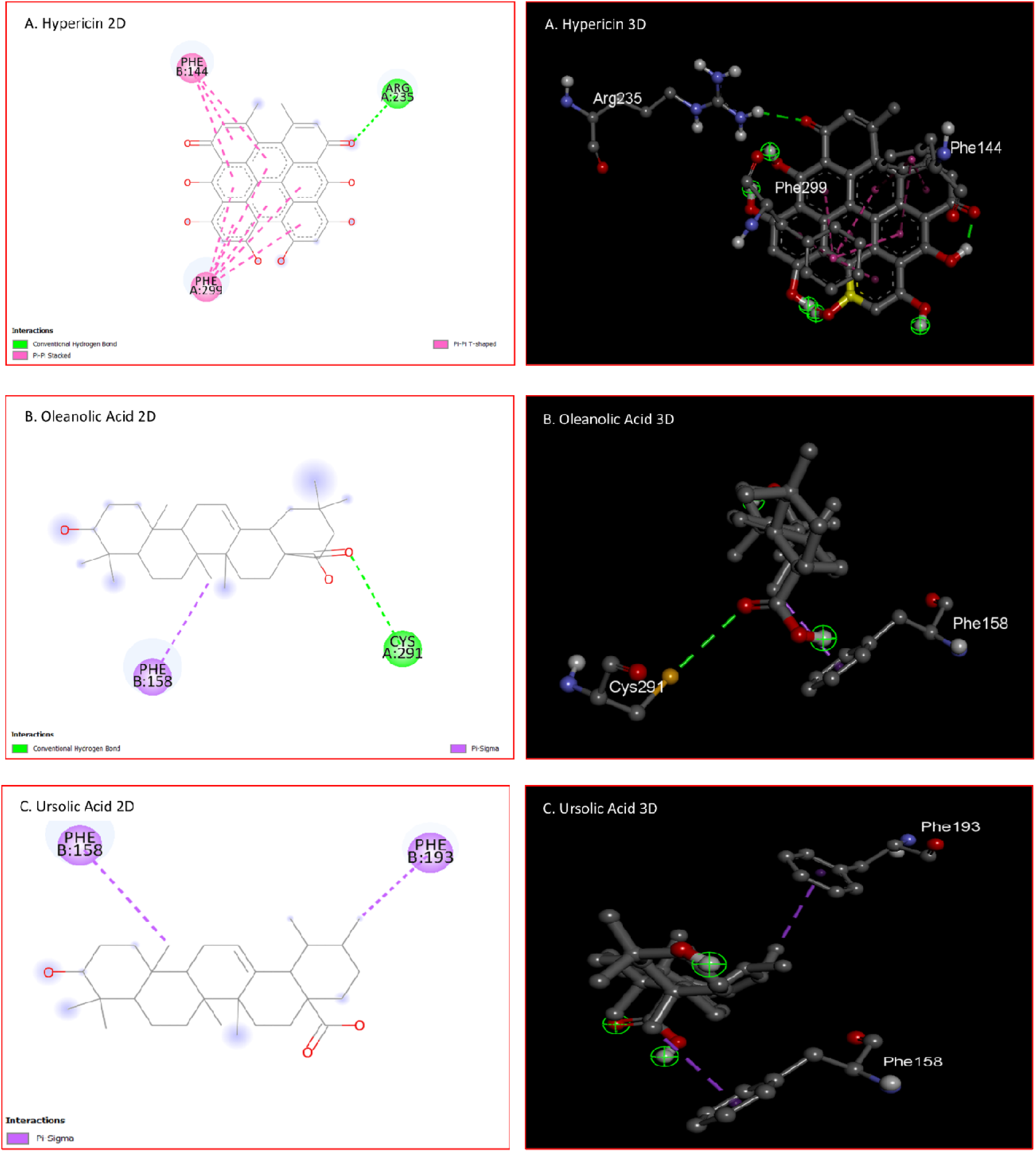
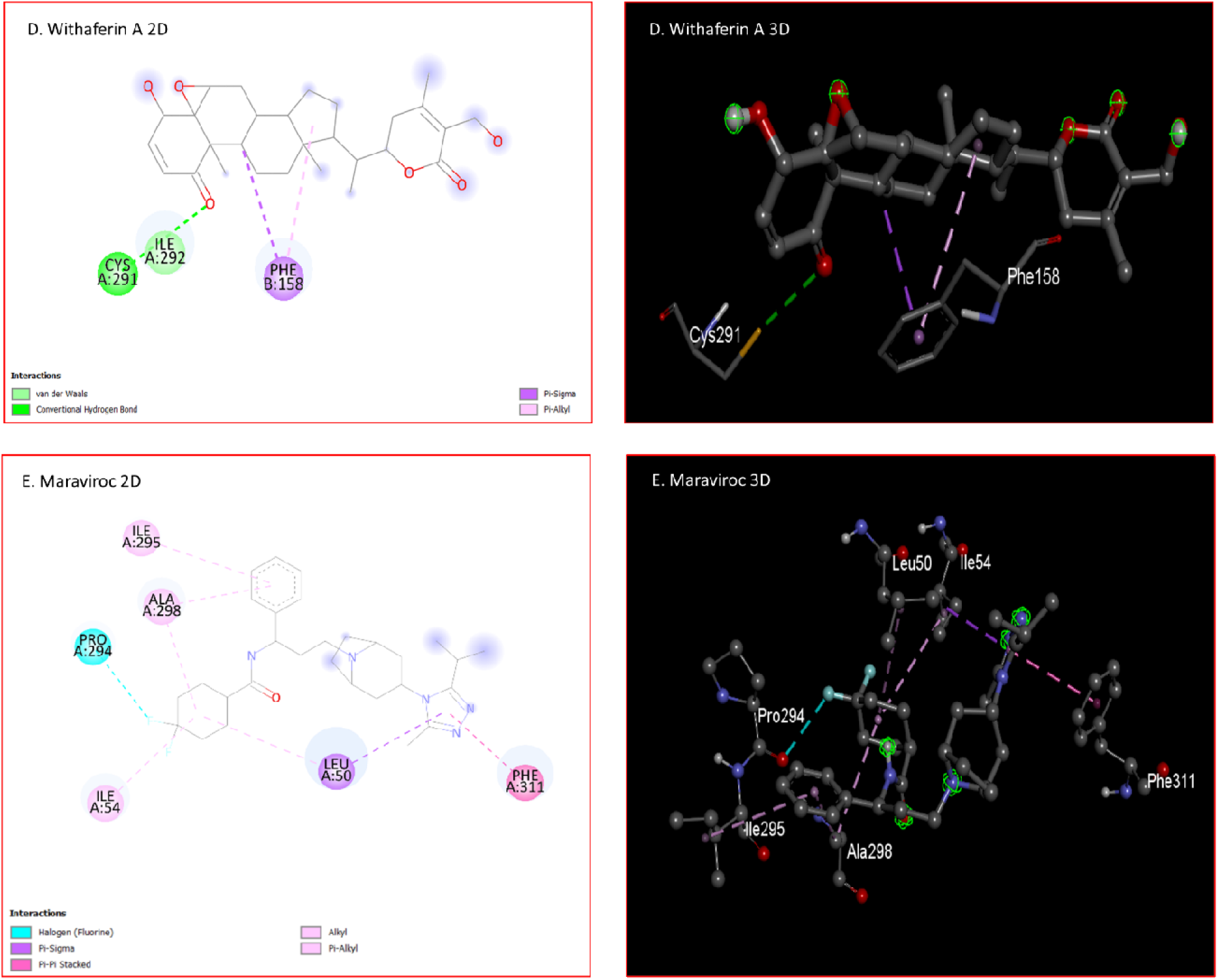
2D and 3D images show the various bonds and amino acids interacting with the five best-selected ligands with the 4MBS receptor (A. Hypericin, B. Oleanolic acid, and C. Ursolic acid, D. Withaferin A, and E. Maraviroc). Interacting amino acid residues of the target molecule are outlined in the diagram, and dotted lines depict the interaction between the ligand and receptor

Finally, the screening was followed by ADMET profiling, druglikeness property, molecular docking, P450 SOM prediction, PASS prediction, molecular dynamic simulation, and selecting the best candidates. Overall, our research suggests that among the selected phytochemicals, withaferin A could be considered as the best performing phytochemical to combat HIV infection by targeting the CCR5 receptor. Withaferin A showed the most consistent result in all the aspects of this research, followed by camptothecin, which also showed quite satisfactory results in many of the sections of our study. However, we suggest further in vivo and in vitro analysis on all the best selected phytochemicals to finally validate the outcome of this study.

**Table 12.** Binding features of top compounds selected against 3ODU and 4MBS.

| S. No. | Target | Name | Hydrogen Bond Interaction | Other Bond Interaction |
| --- | --- | --- | --- | --- |
| 01 | 3ODU | Ursolic Acid |  | Pi-Sigma: TRP125 |
| 02 |  | Oleanolic Acid |  | Pi-Pi Stacked: TRP125<br>Pi-Alkyl: PRO163, VAL206, ALA164 |
| 03 |  | Theaflavin | PRO147, ASN143 | Pi-Cation: HIS140<br>Pi-Sigma: LEU151<br>Pi-Pi Stacked: HIS140<br>Pi-Alkyl: VAL155, LEU132 |
| 04 |  | Hypericin | TYR135, SER224 | Pi-Sigma: LEU150, VAL155, ILE221<br>Pi-Pi T-shaped: TYR135<br>Pi-Alkyl: ILE221, VAL155, LEU150 |
| 05 |  | Camptothecin |  | Pi-Pi Stacked: TRP125<br>Pi-Alkyl: PRO163, VAL206, ALA164 |
| 06 | 4MBS | Hypericin | ARG235 | Pi-Pi Stacked: PHE299<br>Pi-Pi T-shaped: PHE144 |
| 07 |  | Ursolic Acid |  | Pi Sigma: PHE158, PHE193 |
| 08 |  | Withaferin A | CYS291 | Pi-Sigma: PHE158<br>Pi-Alkyl: PHE158 |
| 09 |  | Oleanolic Acid | CYS291 | Pi-Sigma: PHE158 |

**Figure 10 (i).**
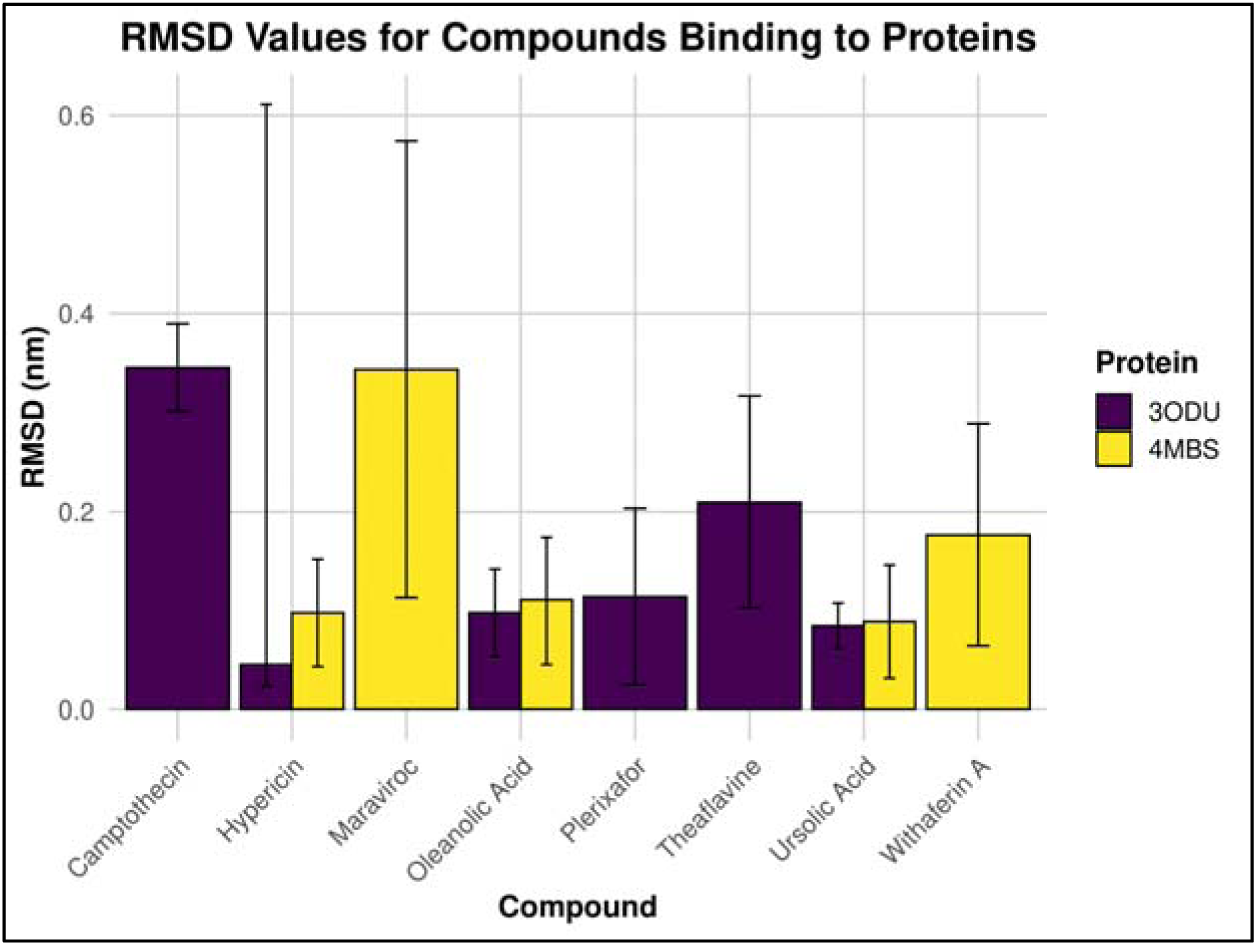
Average RMSD Analysis

**Figure 10 (ii).**
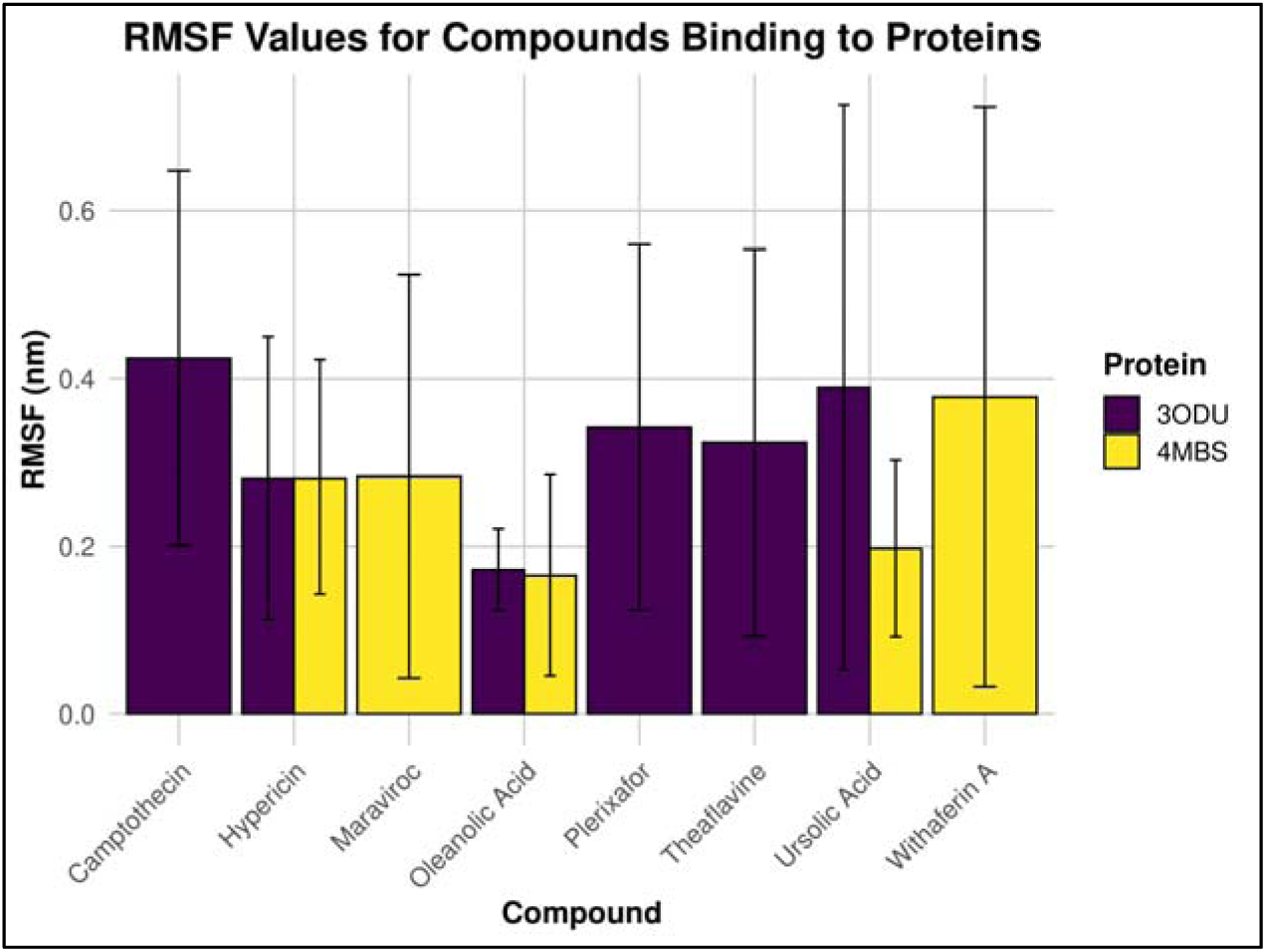
Average RMSF Analysis

**Figure 10 (iii).**
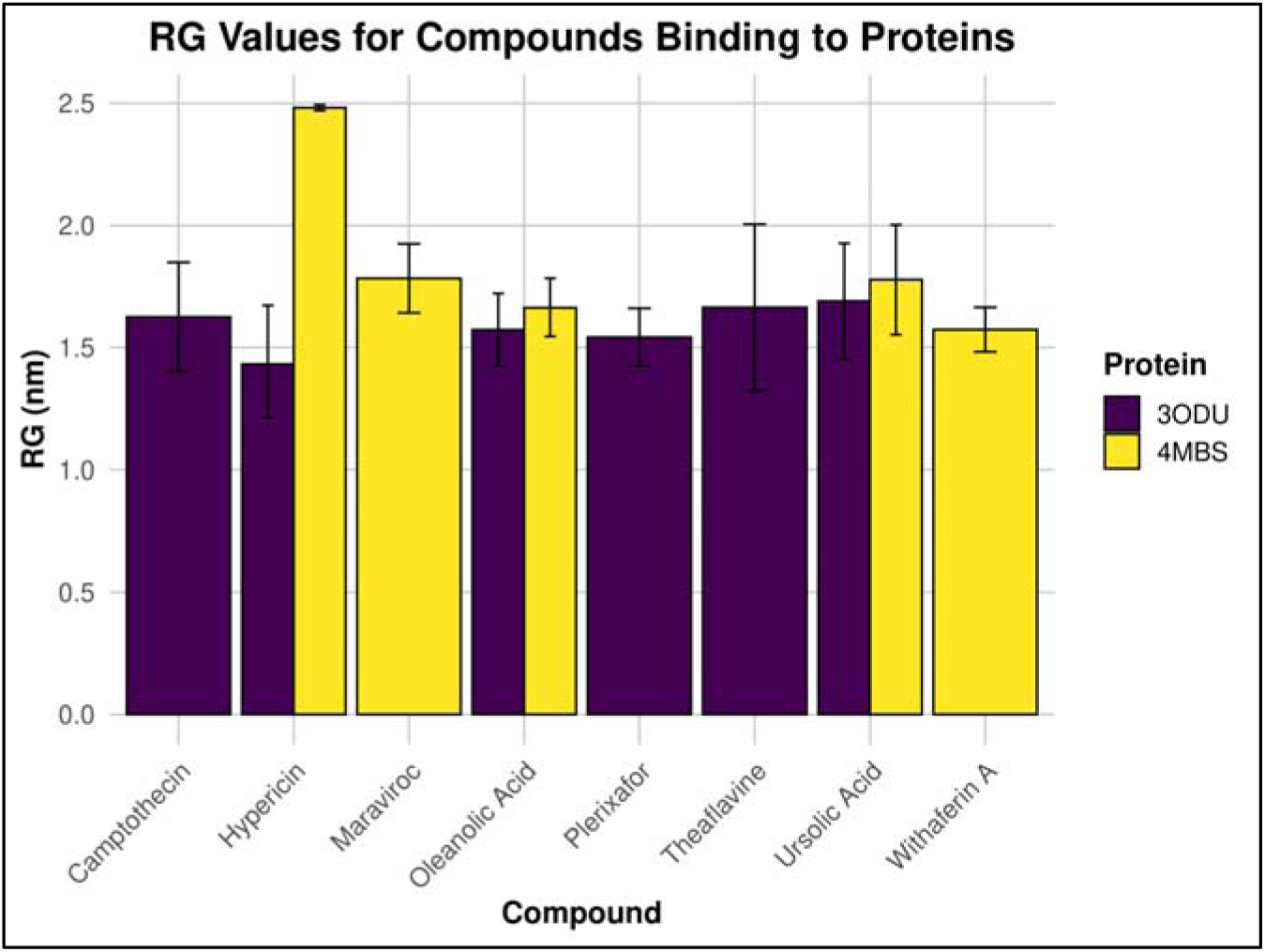
Average Radius Gyration Analysis

**Figure 10 (iv).**
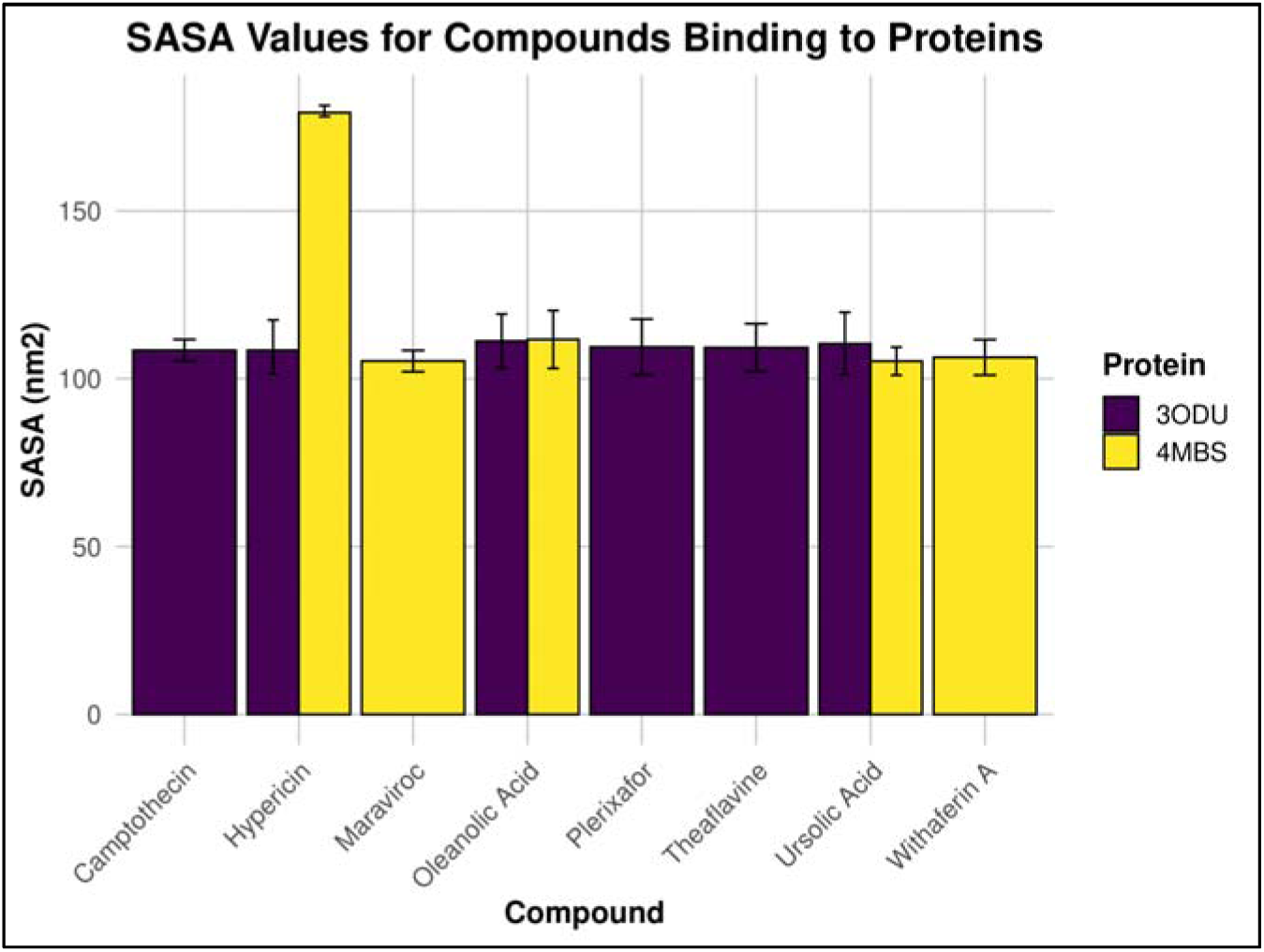
Average SASA Analysis

**Figure 10 (v).**
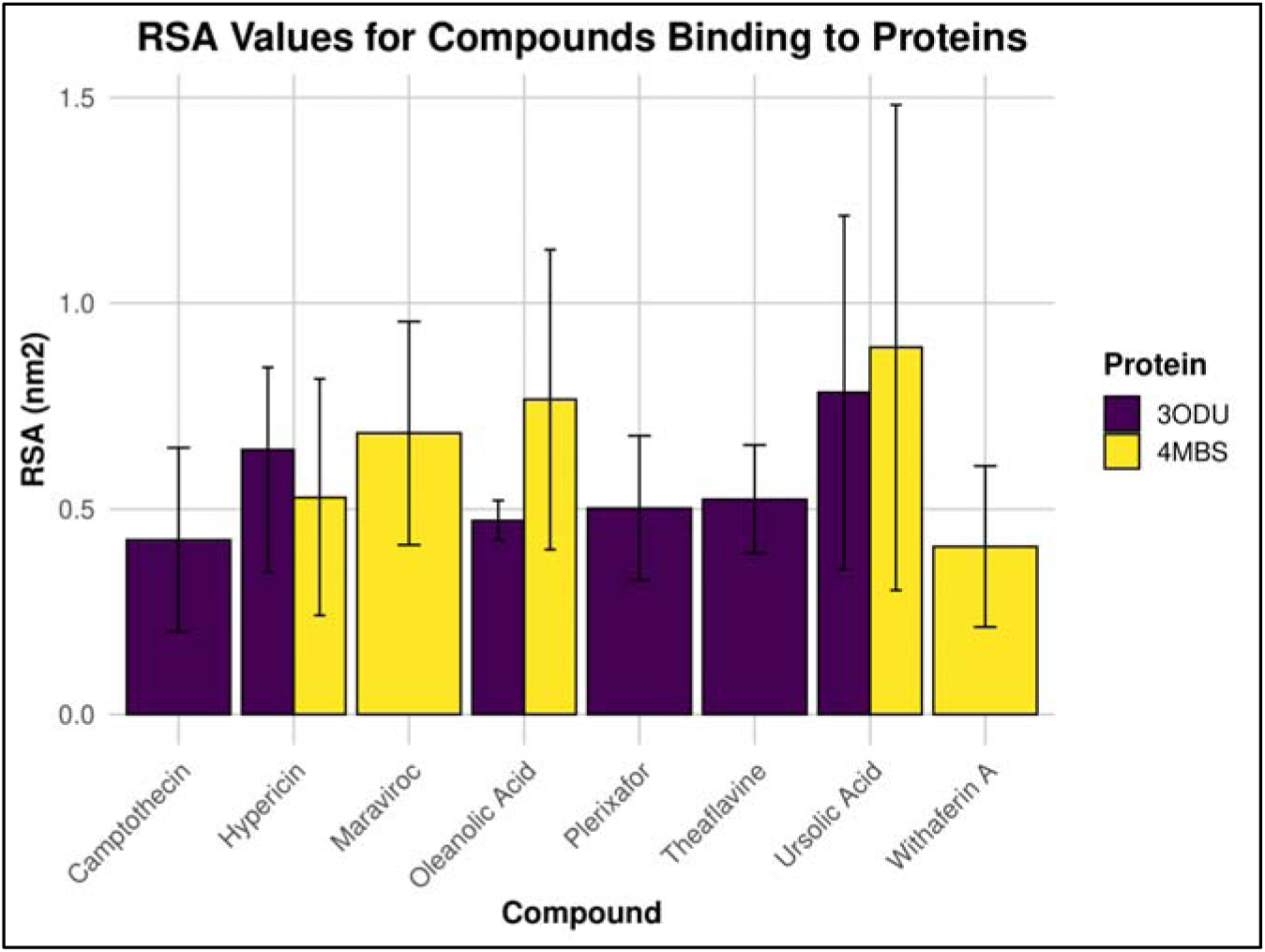
Average Residue Surface Area Analysis

## 5. Conclusion

In the experiment, fifty-three anti-HIV compounds from plants were examined against CCR5 and CXCR4, of three distinct pathways that cause HIV infections. Nine ligands were investigated for each group utilizing various computer-aided drug discovery techniques. Continuous computational experimentation revealed that Withaferin A, Oleanolic Acid, Ursolic Acid, Theaflavine, Camptothecin, and Hypericin were the most effective CCR5 and CXCR5 inhibitors. Then, their drug potential was evaluated in several post-screening trials wherein they were predicted to perform similarly and well. However, the authors recommend that further in vivo and in vitro research using wet lab methods be conducted on the six best and remaining agents to establish their potential, safety, and efficacy.

## 6. Declarations

### Ethics approval and consent to participate

Not applicable.

### Consent for publication

Not applicable.

### Availability of data and material

All the data and material are mentioned and available in the original article.

### Authors’ contributions

SSN, BS, TAA, and RM conceptualized and designed the study; SSN, TAA, and BS conducted the experiments with the help of SM and SSI; BS, SSN, RM, USZ, and MSR prepared the manuscript and RM supervised the study.

## Acknowledgements

We thank Nebir et al. and Ferdousy et al. as we used their manuscripts in preprints as two references in preparing this manuscript (119, 120).

